# Synergistic collagen-condiment: Streptococcal collagen-like (Scl) protein in cell-adhesion and diabetic wound-closure matrix

**DOI:** 10.1101/2021.12.02.470992

**Authors:** Rupneet Singh, Chinmayee Choudhury, Kaniyappan Nambiyar, Swati Sharma, Lakhwinder Singh, Alka Bhatia, Dibyajyoti Banerjee, Ashim Das, Anuradha Chakraborti

## Abstract

Group A streptococcus (GAS), *Streptococcus pyogenes* manifests plethora of diseases through its explicit virulence factors. Among these, the recently deciphered MSCRAMMs, Streptococcal collagen-like (Scls) adhesins are most studied proteins in context of their biophysically stable collagenous-sequence (Gly-X-Y) despite the difference from analogous mammalian-collagen. Based on recent evidence on collagen-mimetic Scls, we elucidated biomaterial-potential of the unmodified, recombinant Scl1 (rScl1). Initially, rScl1 trimeric- assembly yielded its stability *in silico* than the monomeric-unit. Thereby, rScl1 matrix characterization was confirmed *in vitro*. rScl1 exhibited high A549 and HepG2 cell-viability—rScl1 dose incremented to 20.0 µg/ml at time points up to 24 hr, and on 24 hr stored-dishes—deliberating it non-cytotoxic. Imploring cell-adhesion potential, we observed increased cell-counts tangential to rScl1-gradient. This affirmative prelude on rScl1 as a supporting-matrix cued its synergy to collagen; we discerned it through rScl1-augmented, full-thickness diabetic wound-closure *in vivo* and as a first, we studied > 18-month rabbit alloxan-models. We have ascertained re-epithelialization with higher type III collagen in absence of inflammation evidenced morphometrically and histologically. Finally, we correlated our observations through atomistic-evaluation of rScl1-α2β1-integrin interaction, surprisingly, with augmented binding-energy compared to collagen. Hence, connoting recombinant-streptococcal collagen as an ‘alternate’; with further characterization, rScl1 can potentiate important revelations conceding homogeneous and safe, bio-available, biomaterial.

## 1. Introduction

*Streptococcus pyogenes* has a distinct human specific disease-spectra of invasiveness from mild skin-infections to suppurative necrotising fasciitis, to autoimmune sequels which signify its range of virulence-factors. Recent inclusion in the cell-surface virulence factors is the ‘Streptococcal collagen-like’ (Scls)-adhesins. A collagen-like domain is associated with Scls leveraging these proteins the essential pathogenic potential(s) as MSCRAMMs. Previously, these adhesins, Scl1, Scl2, have been studied biophysically, especially towards a physiologically stable, collagen-receptor binding-motif(s) bearing protein [1–3]

As proof of principle, certain molecular-domains with structural and functional identities to collagen have been conserved in many human proteins and other domains of life [4–6]; moreover, collagen-trimer is credited to be among the fundamental protein-structural elements. Recently, bacteria have been found to carry collagen-like sequences, whereby only few express these [7–10]. In our previous-studies, homology and conservation of these genes were recognized with 100% expression in group A streptococcus (GAS)-strains from 27 *emm* types [11]. Our another-study identified M1-2 (*emm1-2* serotype), an M1 variant, emerging in northern India; its sequence and annotation is recently updated [12]. In this study, among the virulent rheumatogenic-M1, the novel M1-variant M1-2, and the nephritogenic-M49 proteins Scl1 and Scl2, we have studied the full-length Scl1-adhesin from serotype M1.

Matrix and its abundant component, collagen with distinct polymeric features manifests importance beyond-structure and provides functional-cues to its microenvironment; yet collagen is the diverse protein-family that shows tremendous conservation. In human collagen, this conservation implies structural features and is evidenced through 1/3^rd^ of X & Y-residues in the -GlyXY- repeats being proline and its hydroxyl-forms [13, 14] yielding stability. Above and beyond, the functional aspect connotes the signalling (distinct receptors), the coordinated-interaction of collagen commencing with the neutrophil activation [15] to terminal-stage healing in keratinocyte, and the myofibroblast mediated wound-matrix reorganization (and contraction) [16]. Thus, being the major component of extracellular matrix (ECM), role of collagen is evident towards a normal wound-healing mechanism. In extracellular milieu, collagen provides cues to the cells as its synthesis increases temporally throughout healing replacing the collagen type III with type I; apparently type I collagen is the abundant form in a healthy cutaneous tissue [17–19].

Notably, towards clinical applications the conventional sources of collagen processing have been suggested to incur concerns on disease-transmission risk, bear product variations and a high cost [20–22]. With immense properties of Scl(s) in consonance to collagen, pertaining to both the structural requisite of triplet-repeats leading into a stable structure, and the functional feature owing to the inherent, receptor binding-motif to collagen; Scl(s) presents bio-availability, non-immune-activity and are a fundamental-polypeptide biomaterial which might augment the ECM-milieu. Constructs with tandem-repeats (Scl2) are reported, as the natural length of their collagenous-domain is only few-100 residues long against the length of human collagen [23]. Although, some studies have focused on chemical-modification in Scl2, it is unknown if it could incur some untoward effects. As a first proof of principle study, we analysed the unmodified full-length molecular-domains of collagen-like protein in full-thickness, cutaneous diabetic-rabbit.

To evaluate biomaterial potential of recombinant streptococcal collagen-like protein (rScl1), we cloned all domains from Scl1-cds except the signal-peptide and the Gram-positive (C-terminus) wall-anchor. We characterized the concentrated-rScl1 in biomaterial context and demonstrated an innocuous and non-cytotoxic, bacterial collagenase-sensitive matrix. We devised and established, *in vitro* rScl1 adherence potential (in cell-culture), additionally, we studied storage of rScl1-coated plates for few days, at 4°C to understand their amenability for prolonged cell-culture studies. With this prelude, we demonstrated *in vivo* wound-closure analysis on the potential of rScl1 protein with manifestation in cutaneous wound in diabetic-models. Recombinant streptococcal collagen-like protein being biocompatible and non-immune-active; it’s unmodified-multidomain soluble formulation seems to synergise the role of inherent, underlying-collagen matrix in diabetic-cutaneous healing. Conclusively, furthering with the *in silico* atomistic-modelling we attempted to confirm stability of the collagen-integrin receptor interactions with variable rScl1-residues modelled as ten ‘integrin-binding motif peptides’ and establish the rationale towards the functional aspect of rScl1 integrin binding. The response of rScl1-peptides with variations pertaining to their specificity towards integrin-α2β1 predicted interactions showing higher binding-energy against collagenous-domain of type I collagen.

The futuristic functional studies of this protein might prove to be an innocuous and novel alternative to collagen, which obviates disease transmission concerns, and rather secondarily increases efficiency of healing; a recombinant molecular approach necessitates their study.

## 2. Materials and methods

### 2.1. Bacterial strains

Group A Streptococci (GAS) serotypes, M1 (type strain; ATCC 700294), M49 and M1-2 (clinical strains) were studied. Strains employed were collected from lab-stocks at Dept. of Experimental Medicine & Biotechnology, PGIMER, Chandigarh. Accession numbers (IDs: GenBank) for the strains are, M1 *scl1*, Scl1: AE004092.2; AAK34668.1; *scl2*, Scl2: AE004092.2; AAK33941.2 and M49 *scl1*, Scl1: CP000829.1; ACI61890.1; *scl2*, Scl2: CP000829.1; ACI61139.1. GAS strains were cultured on Columbia agar supplemented with 5% sheep blood at 37°C, 5% CO_2_.

### 2.2. Cloning

GAS Genomic DNA was extracted by CTAB method [24]. Among virulent, rheumatogenic-M1; the novel M1-variant, M1-2 and nephritogenic-M49 strains, protein domains were analysed *in silico*, selected for desired full-length sequence and primer designing. *scl1*, (*scl1*-CL) amplicons were obtained through PCR, the conditions are appended in Table S1 (see Supplementary file).

*scl*-amplicons were purified and digested with Fast-digest restriction enzymes (NEB) for directional cloning; single digests (BamHI and NdeI) were analysed on 1% agarose gel for complete digestion, gel purified and processed for sequential double-digestion (Sal1 and XhoI). Similarly, the pQE30 (Qiagen, Germany) & pET28a vector-DNA (Novagene) were digested and purified. Ligation was achieved using T4 DNA Ligase (Promega, USA) with incubation at 4°C for ∼16 hours. *E. coli* DH5α, cloning host, competent-cells prepared using CaCl_2_ method were transformed. The potential positive transformants [DH5α (pET28a:*scl1,* pQE30:CL)] were replica-plated on LB agar plate containing Ampicillin 100 µg/ml and/or Kanamycin 40 µg/ml. Screening was done by colony-PCR using both insert- and vector-specific primers. Standard Maniatis protocols [25] and manufacturer’s instructions were employed as needed. The schematic for the *scl*-constructs referred in this study are shown in Fig. S1A-C, (see Supplementary file).

### 2.3. Soluble pRS-1

#### 2.3.1. Over-expression of fusion/recombinant Streptococcal collagen-like protein 1 (pRS-1)

The construct after screening and sequence-confirmation of positive transformants (commercially) was selected for Scl1 over-expression in *E. coli* BL21 (DE3), achieving induction with 0.5; 1.0 mM IPTG at 37°C/4 hrs. The soluble- and cell-debris fractions were checked (for over-expression) by Coomassie staining, Silver-stain on 10% SDS-PAGE and further confirmed through immuno-blotting on PVDF membrane using anti-His primary antibody. 500 ml and 1 L cultures were optimized and protein yields were analysed. Overnight activated-culture as 1% starter-culture was used, cultures were shaken obliquely, at 200 rpm until O.D. of 0.4 to 0.5 was achieved which proved best for IPTG induction for rScl1. Kanamycin as selection antibiotic (35 µg/ml) was supplemented as required.

#### 2.3.2. Purification of the recombinant Scl1 (pRS-1)

Recombinant-construct with N-terminal His-tag was purified through IMAC-based Ni-NTA (Qiagen). IPTG induced, over-expressed bulk-culture was pelleted, lysed by sonication (30 sec On/Off cycles) and lysate harvested as the bulk soluble-protein in cleared lysate. Resin was incubated in cleared-lysate (30 min., on rotator) after equilibration (50 mM NaH_2_PO_4_, 300 mM NaCl (pH 8.0)). Subsequently, the resin was packed into a column and purified using imidazole-gradients. Elutes were collected and checked on 10% SDS-PAGE. Concentrated protein was achieved using Millipore protein-concentrator spin-columns after desalting through pre-packed PD10 columns [GE]. All steps were performed on ice.

### 2.4. Characterization of the recombinant Scl1 (pRS-1)

#### 2.4.1. In vitro Integrity and in silico homology of the recombinant Scl1 (pRS-1)

Desalted purified protein was simultaneously buffer-exchanged into three different buffer-systems; Bicarbonate carbonate (0.1 M, pH 9.6), Phosphate buffered saline (pH 7.4), Tris (0.1 M, pH 8.0). It was characterised for solubility, stability and integrity over storage in these buffers after concentration, (separately in all buffers) through immunoblotting using anti-His primary antibody raised in mice and monoclonal (goat) anti-mouse secondary HRP-conjugated antibody (Sigma). Further, three different estimation-methods were employed towards a correlation, viz. UV absorption-spectroscopy, Bradford- and Bicinchoninic acid (BCA)-estimations. *In silico* homology analysis between recombinant rScl1 CL-domain and human type I α chain [position 162-1218 without N- and C-terminal pro-peptide domains] using protein BLAST and evaluation for features of sequence-residues lending stability using Expassy Protparam tool, was done.

#### 2.4.2. Collagenase sensitivity of recombinant protein (pRS-1)

Purified, concentrated pRS-1 was further characterized for sensitivity to the action of bacterial-collagenase, (*Clostridium histolyticum,* Sigma). Crude, 300 µg/ml and 1:1 diluted- pRS-1, 150 µg/ml, solubilized in PBS buffer (0.1 M, pH-7.4), were exposed to 0.054 U, 0.108 U and 0.162 U of bacterial-collagenase for 1 min. The assay was performed in 96 well-plates, in triplicates, incubated at 37°C followed by freezing the plate to inhibit the enzyme. Extent of collagenase-sensitivity was achieved by pRS-1 integrity analysis by 10% SDS-PAGE followed by Coomassie staining.

#### 2.4.3. Cell-culture matrix analysis of the purified-recombinant protein (pRS-1)

Briefly, 96 well-plates coated for 2-3 hrs (37°C) or overnight (4°C) with pRS-1 gradient- 0.5, 2.5, 5.0, 7.5, 10.0 and 20.0 µg/ml, were aseptically stored. Coating in bicarbonate carbonate (0.1 M and/or 0.3 M, pH 9.6), blocking (5% BSA) and washing with TBST (0.05%)-3x and TBS-1x were done. Subsequently, the coated-pRS-1 was probed using anti His-primary and anti-mouse HRP-conjugated secondary antibody, respectively. HRP substrate-TMB colourimetric detection was done after 15 min. incubation at 37°C at 620 nm (BioPhotometer, Eppendorf, Germany). Coated plates were stored (until 24 hrs) packed with parafilm at 4°C and used as required.

### 2. 5. Cytotoxicity assessment of purified-recombinant protein (pRS-1)

Hepatoblastoma HepG2 cells and pulmonary epithelial A549 cells, cultured on pRS-1- coated 96-well plates were assayed (for any cytotoxicity) using MTT (Sigma) cell-viability method, in triplicates, for both cells, assessed at 12 hrs, 24 hrs. MTT [3-(4 5-dimethylthiazol- 2-yl)-2 5-diphenyltetrazolium bromide] assay was done as per standard protocol [26]. Briefly, about 5000-7000 cells were seeded per well, incubated at 37°C, 5% CO2; HepG2 cells were cultured in MEM (Sigma) & A549 cells in RPMI 1640 (Sigma), FBS supplemented. Cells were cultured to near-confluence (in 96 well plates), pre-coated with 0.5, 2.5, 5.0, 7.5, 10.0 and 20.0 µg/ml pRS-1. After apt incubation, the monolayer was washed (PBS pH 7.4, 3x) and MTT solution (1 mg/ml) was added along with appropriate media. The plates were incubated at 37°C, 5% CO₂ for 4 hrs, then treated with the solubilization agent, dimethyl sulfoxide (DMSO). Reaction was done on rotating platform (15 min./RT) to ensure complete solubilization of blue formazan-product, read at 570 nm. Results were analysed taking media-only as blank and cells cultured in uncoated-wells as controls.

### 2.6. Cell-adhesion analysis of recombinant protein (pRS-1) in vitro

Briefly, HepG2; A549 cells were seeded in 96 well-plate and monitored for their optimum morphology on coated-pRS-1 gradient (0.5, 2.5, 5.0, 7.5, 10.0 µg/ml and 20.0 µg/ml), in triplicates. Morphology was observed microscopically, upto 24 hrs. Cells were fed with 10% FBS in apt media.

### 2.7. Analysis of recombinant protein (pRS-1) in cutaneous wound healing in experimental- animal model (in vivo)

New Zealand white rabbits or *Oryctolagus cuniculus* (8-10 weeks, body weight 1.7-2.0 kg) (male and female) were developed as experimental-diabetic models. All animals were fed on standardized pellet diet, vegetables; and, water was accessible ad libitum. They were housed and maintained in 12 hr-light/dark cycle in controlled-temperature and humidity at the Small Animal House, PGIMER, Chd. Animal studies were conducted in accordance to the PGIMER, Chd/Institute animal ethics committee guidelines (IAEC).

#### 2.7.1. Alloxan induced experimental-diabetic rabbit model

Diabetic-model was prepared through alloxan monohydrate (Sigma Aldrich), single and double-administration. Alloxan preparation was done in saline (sterile inj.) on ice and was administered via marginal-ear vein (rate-1.5 ml/minute). Post administration, phased- response of hyperglycemia was precisely followed (glucose-test strips; DrMorepens). The first 24-48 hr critical-phase, which induces hypoglycemic-shock was optimized; animals were closely monitored along with regular subcutaneous-injections (dextrose 5-10 %). Besides, after treatment, to avoid possible hypoglycemic-shock, glucose was given orally until 48-hr, at 2, 4, 6, and 8 hours and so on (for 2 days post allox. I.V.). Food and water intake were monitored daily. Insulin-treatment was not employed in any animal. Hyperglycemia was developed after 48 h and stabilized after one-week. The blood-glucose level more than 250 mg/dl was considered hyperglycemic [27]. Dose and all optimized observations are appended in Table S2, (see Supplementary file).

#### 2.7.2. Rabbit ear-ulcer model development: Wound bed generation

Under local anaesthesia, Lignocaine (2% inj. subcut.) wound beds were created in the ears. Briefly, all hairs were removed from ear-pinna and punch-biopsy was employed, 2.5 mm; 6.0 mm sterile tool. Full-thickness wound-beds with secure cartilage underneath were carefully made [28]; sterile conditions were ensured. After formation of wounds, analgesics (Tremadol Hcl inj. subcut.) were given with regular cleaning of diabetic-models. Grouped distribution of animals is appended in Table S3, (see Supplementary file). Animals were grouped as controls/untreated groups- **control-healthy** (group-I); **control-diabetic** (group-II); and test groups- the pRS-1-**treated-healthy** (group-III); pRS-1-**treated-diabetic** (group-IV) and the positive control-treated with commercial ECM (Matrigel, Sigma) diabetic-models (group-V).

#### 2.7.3. pRS-1 augmentation and synergistic wound closure analysis

Control- and diabetic-models were treated with purified-pRS-1 at optimized concentration (10 µg/ml-2.5 mm & 20 µg/ml-6.0 mm wound-beds, approx.). Tegaderm^TM^ (U.S.) was used to cover treated wound-beds as needed. Models were monitored and observations recorded day-wise until wound-closure. Wounds were extracted after 15-20 days (after sacrificing models), stored immediately in normal buffered formalin and processed. Tissue transverse-sections were studied using three distinct stains; H&E, Gomori and Picro Sirius red.

Battery of assessment-tools used for wound-closure included wound-bed image gross analysis; morphometry of images and tissue-sections using NIS Elements software (Nikon, Japan) and ImageJ. Parameters *viz.* closed wound-area and cross-sectional wound-bed length etc. over days were monitored.

### 2.8. Statistical analysis

The GraphPad Prism V. 7 (Inc. CA, USA) was employed for the statistical analysis. One-way and Two-way ANOVA with multiple comparisons was made as required. Results were expressed considering ‘p value’ less than 0.05 as significant.

### 2.9. Computational Modelling of rScl1 (pRS-1) structure and it’s interaction with human integrin

#### 2.9.1 Modelling of the 3D structure of Scl1 protein

The FASTA sequence of M1 GAS Scl1 (*scl1*) was retrieved from UniProt [29]. An initial BLAST search [30] with the sequence returned 3HVQ (Molecular Envelope Structure of Type I Collagen) as the best hit with 40% local (Fig. S2A) and 38% global (Fig. S2B)sequence identities (calculated using Clustal Omega [31]), respectively. However, it was not possible to build a structure from 3HVQ as this structure had a very low resolution with only the coordinates of Cα atoms resolved. This sequence similarity gave a clue about the overall structure of the collagen-like domain. So, the structure was initially modelled in two parts. The N-terminal residues 7-73 were modelled (model1) using the Modeller [32] considering the crystal structure of the globular domain of M3-type group A Streptococcus Scl2 (*scl2*) as template. The sequence identity between Scl1; Scl2 were as low as 22%. So after homology modelling, several iterations of energy minimizations and structure quality check with the ProCheck [33] and ERRAT [34] servers were carried out in order to refine the structure. Finally, a reasonably good model was obtained with overall ERRAT quality score of 92 (Fig. S3A). ProCheck analysis showed 93.5% of residues in the allowed regions and only one residue in the disallowed region, ensuring a good model (Fig. S3B). In absence of a homologous template structure for the residues 74 to 279, the FASTA sequence for this region was submitted for *ab initio* modelling using I-TASSER server [35]. As, I-TASSER can process only 200 residues at a time, residues 80-279 were submitted. I-TASSER returned 5 best models for residues 76-279, with c-score ranging from -3.05 to -4.48. The third model with c-score -3.90 (model2) was chosen for our study considering its collagen-like helical linear structure resembling 3HVQ. ERRAT quality factor for this model was 97.36 (Fig. S3C), which is considered as a fairly acceptable model. Then, a chimeric structure for the full Scl1 protein was built by merging model1 and model2 using Modeller. Residues 74-79 were added as a loop between model1 and model2. This monomeric model structure was then submitted to the SymmDock server [36] to build a trimetric structure using symmetric docking. SymmDock returned top 10 trimeric solutions for Scl1. The top scoring solution with a globular N-terminal alpha helical head (similar to Scl2 protein crystal structure) and a collagen-like tail was selected for further study (Fig. S4A).

#### 2.9.2 Generation of Integrin-Scl1 peptide complexes

In order to study the affinity of Scl1 with human-integrin we considered the 2.1 Å resolution crystal structure of human-integrin (1DZI) complexed with a 21 residue stretch of human collagen triple helix. The CL-domain of Scl1 (from G69 to N257; GenBank: AE004092.2; AAK34668.1) was divided into nine consecutive 21-residue and one 22 residue peptides (Table 1).

**Table 1.**
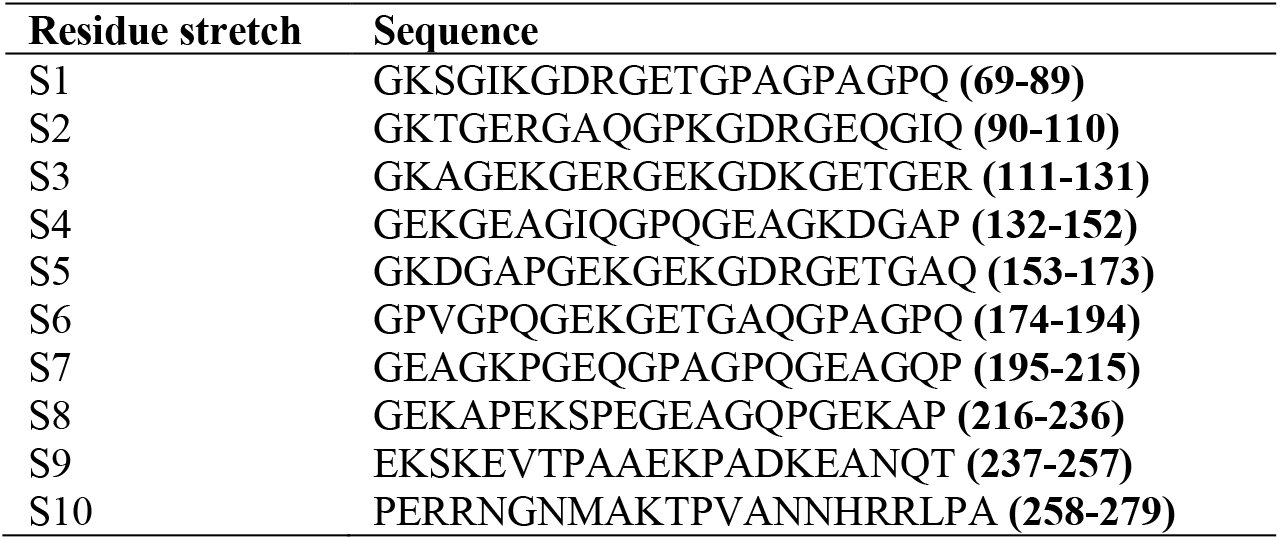
Details of the 21 residue peptides extracted from the collagen-like domain of Scl1 to study their interactions with human-integrin

Each residue of the human collagen trimer in its complex with 1DZI was systematically mutated to obtain the new complexes with 10 Scl1-peptides. Each residue in the collagen trimer-1DZI complex was changed to convert it to Scl1 peptides. Thus, we obtained 10 complexes of human-integrin with 10 different 21-residue Scl1-peptides (trimeric form). The 10 integrin-Scl1 peptide complexes along with the integrin-human collagen complex were energy minimized under OPLS3 force field and VSGB solvation model with dielectric constant 80.00 and convergence gradient 0.01 kcal/mol using Schrodinger Suite (2019 release).

The complexes were then submitted to PPCheck [37] server for prediction of the binding energies of the human collagen and Scl1-peptides with human-integrin. PPCheck calculated the binding energies in kJ/mol, which we converted to kcal/mol. The H-bonds and other interactions between Scl1-peptides/human collagen was analysed using Maestro interface of Schrodinger suite.

## 3. Results

### 3.1. Recombinant constructs of Streptococcal collagen-like protein, Scls

Scls (proteins) are among cardinal adhesins recently identified in genomes of most GAS-strains and are highly conserved, show distinct interaction with human-major collagens [11], mediated through specific mechanisms [38–40]. In context of these observations, we hypothesized/speculated to characterize Scls from GAS-strains with varied virulence towards similarities, with emphasis on their collagenous-domains. To achieve this, in current study, 2-rheumatogenic and 1-nephritogenic GAS-strains were included. Scl-1; -2 protein-homology and alignment studies between M1 & M49 strains determined complementarity, depicting very slight variations (see Supplementary, Fig. S5A(i); (ii)). Further, the protein-family database study using Interpro-software, revealed similarity in domain-structures for these adhesins, in these strains (see Supplementary, Fig. S5B). In this study, based on the sequence-studies, GAS M1, Scl1 major-domains were selected for full-length and CL(-only) domain recombinant-constructs (see Supplementary, Fig. S1A-C).

The full-length construct constituted of N-terminus V-domain followed by CL-domain and L-domain. Towards over-expression and purification, the positive-clones with full-length, 813 bp, Scl1-domains (Fig. 1A), and 339 bp, small-CL domain (Fig. 1B) were selected after screening and sequence-confirmation. Recombinants ensued a soluble-protein under strong pET-expression. pRS-1 has an observed-mass of ∼50 kDa (Fig. 1C). Apparently, the protein-mass was found to be higher than the predicted-mass of 30 kDa (as calculated from primary construct-sequence). Further, towards direct applications and to analyse stability of pRS-1 in different buffers, purified-protein was buffer-exchanged and concentrated separately in 3 buffer systems; i.e., PBS, Tris and bicarbonate carbonate (B/C) buffers (which were selected for their physiological importance).

**Fig. 1.**
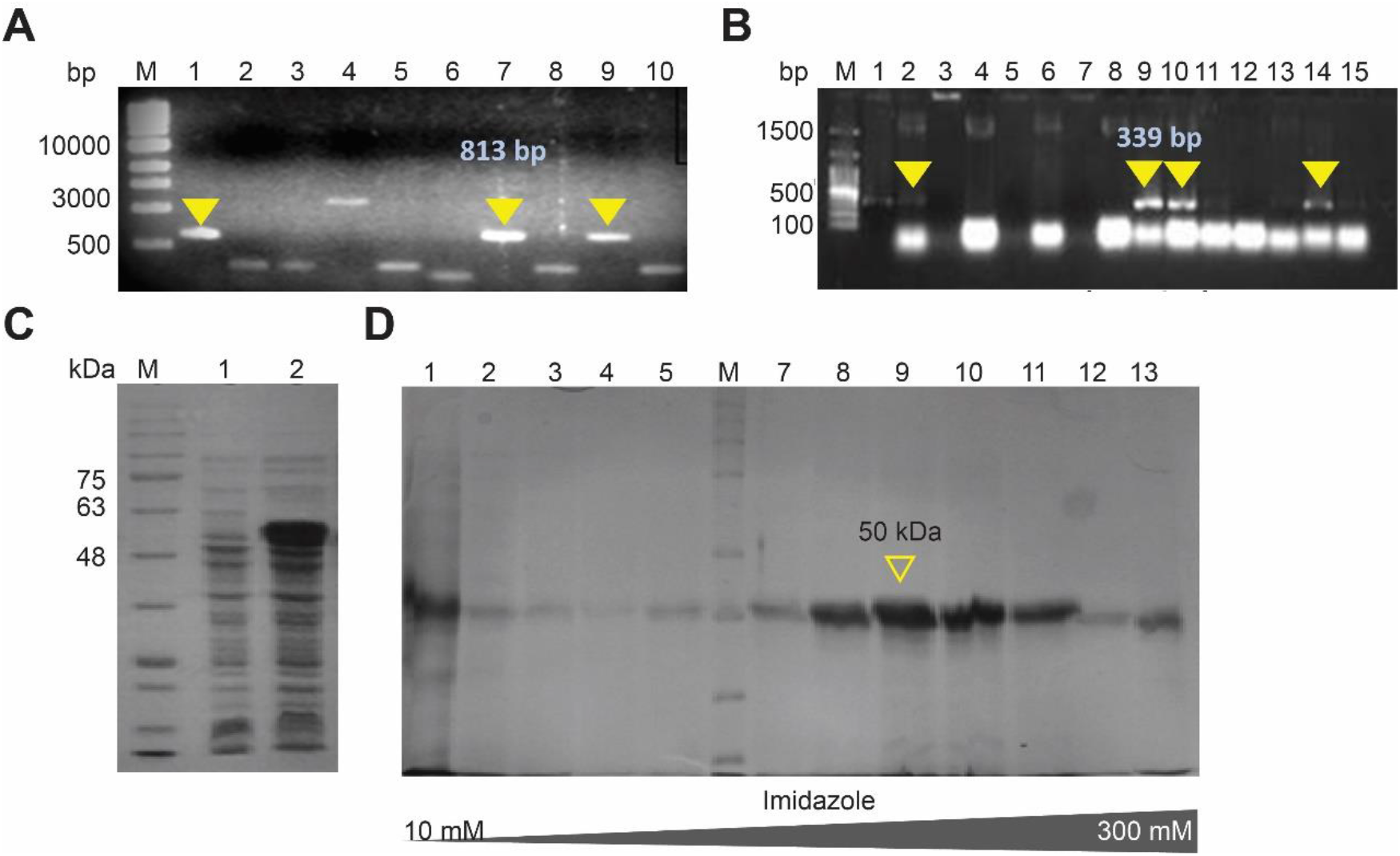
Screening of the positive GAS M1 scl1 clones. (A) 813 bp pRS-1 (rScl1) positive recombinants (Lanes: 2, 4 & 10), M: 1 kb DNA Ladder. (B) 339 bp pRS-CL(rCL) positive recombinants (Lanes: 9, 10 & 14); Lanes 1: (+), M: 100 bp Ladder. Characterization of pRS-1 (recombinant Scl1). (C) SDS- PAGE (10%) showing (i) Over-expression and purification, IPTG (0.1 mM) induction. Lanes; 1: Control (-); 2: Induction (+) 37°C, 4hrs; M: protein marker. (D) pRS-1 purification through IMAC. Lanes; 1: Cleared Lysate [No Imidazole]; 2 to 5: Wash I, II, III, IV [Imidazole 10 mM to 60 mM]; 7 to 13: Purified Elutes [Imidazole: 300mM], M: protein marker.

### 3.2. Characterization lends an easily-expressed, soluble-protein (pRS-1), with homology to human collagen type 1: Structure modelling studies, matrix-like properties and collagenase-sensitivity

#### 3.2.1. In silico analysis revealed 50% homology in pRS-1 Collagen-like domain and human collagen

The EXPASY Protparam tool *in silico* analysis (https://web.expasy.org/cgi-bin/protparam/protparam) revealed the amino acid residue-proportion in pRS-1 backbone that predicted pRS-1-construct stable (see Supplementary Fig. S6A). Further, the collagen-like (CL) domain annotated sequence of GAS M1 Scl1 revealed a > 50% sequence homology (see Supplementary Fig. S6B) to the human collagen type I sequence (leaving aside the tropocollagen terminal domains).

#### 3.2.2. Structural model of rScl1 shows a stable trimeric form and favourable interactions with Human-Integrin

The globular alpha helical domain (residues 7-73) of the trimeric structure is stabilized by inter-chain hydrogen bonds (H-bonds) and salt bridges (Figure 2C). The side chain of W39 of each chain forms an aromatic H-bond with the side chain of Q50 of the adjacent chain. A symmetric and cyclic interaction pattern i.e. W39A-Q50B, W39B-Q50C and W39C-Q50A was observed. Similarly, a salt bridge interaction network was observed between the side chains of K70 and E18 of the adjacent chains in a cyclic manner i.e., K70A-E18B, K70B-E18C and K70C-E18A. Such inter-chain H-bond networks are responsible to stabilize the trimetric form at the N-terminal end. The CL-region comprises several continuous GXY repeats, where X and Y are mostly charged residues and have opposite charges. The oppositely charged X-Y residues of GXY-motif form salt bridge/H-bonds which contributes to the collagen-like linear helical structure of each monomeric chain (Fig. S4C). Also, the polar side chains of X-Y residues of one monomeric chain make H-bond/salt bridge interactions with the main chain of the adjacent chains. For example, if we consider the motif G75-D76-R77 (Figure 2D), the two side chain oxygen atoms D76:A act as acceptors to make H-bond with the main chain –NH groups of D76:B and R77:A. Similarly, D76:B forms H-bonds with D76:C and R77:B; and D76:C forms H-bonds with D76:A and R77:C, which forms a cyclic H-bond network for stabilization of the trimer. Also, the side chain – NH2/NH3+ group of R77A acts as a donor to form H-bond with the main chain carbonyl oxygen of G78A of the next triplet to stabilize the monomeric collagen-like structure. In the C-terminal region, GX-triplets are not found, but the strong cyclic H-bond network among the monomeric chains maintain the trimeric structure. For example, in the C-terminal residue stretch H274-R275-R276 (Fig. S4D), the helical monomeric structure was maintained due to the aromatic H-bonds between H274 side chain and R275 main chain and between H274 main chain and R276 main chain of each monomer. The cyclic inter chain H-bond network among R275:A main chain and R275:B side chain; R275:B main chain and R275:C side chain; and R275:C main chain and R275:A side chain hold the trimeric structure at the C-terminal end.

**Fig. 2.**
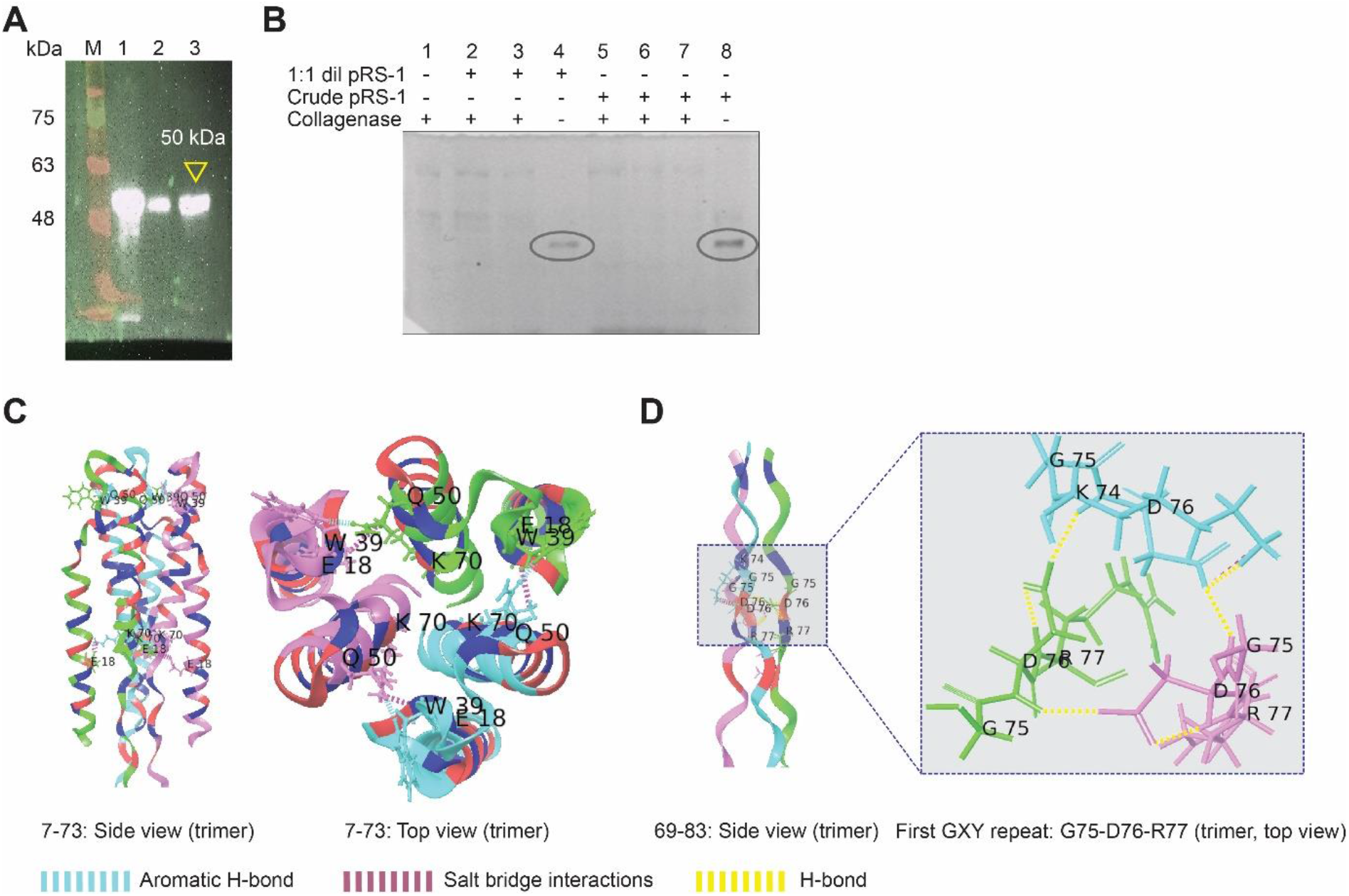
Characterization of pRS-1. (A) pRS-1 Immunoblot (concentrated in different buffers) analysed for N-terminal integrity; B/C (Lane 1); PBS (Lane 2 & 3). (B) SDS-PAGE (10%) showing Collagenase sensitivity (in Units). Lanes: Enzyme (1); Treated (2, 3) & Untreated (4) 1:1 dilute-pRS- 1; Treated (5-7) & Untreated (8) Concentrated-pRS-1. (C) Example of inter-chain interaction pattern in the globular region. (D) Example of inter-chain interaction pattern in the collagen-like domain.

#### 3.2.3. Estimation of total soluble-protein with conspicuous effect of absence of significant amino-acid residues/Intricate protein quantitation and total soluble-protein estimation

*In silico* sequence-characterization of recombinant-construct revealed ‘scanty’ conjugate amino acid (proportion) (see Supplementary Fig. S6A), apparently, the A_280_ based UV-spectra-estimation for pRS-1 demonstrated ambiguous-concentration. We observed, this method demonstrated higher protein amount (and did not correlate with other estimation methods). Besides the absorbance method, pRS-1 was undetectable through Bradford estimation. In this context, total soluble-protein estimation through bicinchoninic acid assay (BCA), which estimates peptide-bonds, proved to be a means of direct quantitation. Further, it was amenable to study concentrated-pRS-1 in different buffers, *viz.*, PBS, Tris and B/C buffers, yielding negligible background. As ‘native-trimer’ is reported trypsin-resistant, the efficient analysis of the construct using MALDI was ineffectual. Thus, resorting to BCA-quantitation (of pRS-1), we found ∼300 µg/ml protein-yield from the pRS-1 batch-buffer-exchange and concentration after IMAC/affinity purification.

#### 3.2.4. Soluble protein, pRS-1 construct was most stable in bicarbonate-carbonate buffer

The 3 low-molar composition buffer-systems studied, PBS, Tris and B/C, were selected to increase our understanding on the solubility and stability of the recombinant protein. pRS-1 at all stages of purification, i.e., after (PBS, Tris and B/C) buffer-exchange and concentration, was checked for integrity and observed-mass through SDS-PAGE. The N-terminal integrity of the soluble-pRS-1 detected by intact N terminal His-tag after 3x-concentration and storage (at -20°C) insinuates a stable behaviour in Bicarbonate-carbonate buffer however, there was loss of protein in PBS on storage, as shown in lanes 2; 3 in (anti-His) immunoblot (Fig. 2A).

#### 3.2.5. Recombinant construct of Streptococcal collagen-like protein depicts collagenase sensitivity

We found rScl1 stable towards trypsin which connotes that it follows the inherent inertness of the trimer. We deliberated purified, concentrated rScl1-construct to be amenable to bacterial-collagenase, as seen in lanes 2, 3 & 5-7 (Fig. 2B), thereafter, we detected its sensitivity on treatment with Clostridial-collagenase.

### 3.3. Coating and cell-viability revealed the recombinant-construct an innocuous cell-culture matrix

#### 3.3.1. pRS-1 coating-studies and visualizing the coated-protein

In our attempts towards matrix-characterization of pRS-1-protein, primarily, the most potential-bio-active material characteristic, i.e., manifestation of augmentation in cell-adherence phenotype, beckoned our studies. We analysed our hypothesis through series of estimations, and at the outset, ensuring direct visualization of pRS-l after coating (in cell-culture plates) was essential (Fig. S7A, see Supplementary). Apparently, direct visualization was imperative but no method permitted to it. Finally, thorough an indirect assay, the coated- pRS-1 was visualized. Using anti-His antibody, detection of pRS-1 N-terminal His-tag was eventually done as a qualitative-indirect assay for the amount actually-coated (in the wells) (Fig. S7C, see Supplementary). Through this approach, we could detect as low as 0.5 µg/ml pRS-1, thus sustainably, we analysed our coated-protein.

Further, left-over buffer after pRS-1-coating was recovered and assayed. We found the left-over buffer free of any traces of pRS-1 as it revealed no spent-protein (Fig. S7B, see Supplementary). His-tag rendered a strong and stable marker here and has been maintained in purified-construct for further use.

#### 3.3.2. Coated-pRS-1 plates assayed for cell-viability and cell-adhesion

Interestingly, in 96 well pRS-1 coated-plates, matrix colorimetric-assay developed indistinct/uniform blue-colour [Blue coloured HRP-TMB oxidation-product in immunolabelling by anti-His antibody] over the gradient ranging 0.5-20.0 µg/ml (Fig. S7C, see Supplementary). We confirmed that the applied pRS-1 was coated each well entirely, as the left-over buffer (after coating) was clear of any spent-pRS-1. Notably, the plates were saturated/set over a period of 2 hrs (37 °C) and/or additionally, over-night at 4°C, before the assay. Prompted to assess the matrix-characteristic, we evaluated the pRS-1 matrix- hypothesis through cell-culture studies (on these coated-plates). We assessed pRS-1 coated-plates for their compatibility in cell-culture and cell-cytotoxicity over a gradient, in 3-setups. Cells cultured for (a period of) 12 and 24 hrs on 2 hr-coated plates depict the first- & second- set, respectively. Subsequently, for the third-set were cells in (over-night) stored-coated plates. We elucidated effects of pRS-1 matrix on cell-adhesion in two distinct growth- efficiency adherent-cells; A549, lung-epithelial; and, HepG2, hepatoblastoma; the cells were not serum-starved. We monitored cells on a time-based manner, observed their adherence, microscopically and their cell-viability using MTT-assay.

Along pRS-1 incremented-gradient (in wells), we observed an increased cell-viability, and normal morphology in adhered-cells, depicting no cyto-toxic manifestations. Specifically, in both A549; HepG2 cells, we observed >70-80% confluence, with percent-viability ranging 80-100% at 12 hr (Fig. S8A and D); and 24 hr culture (Fig. S8B and E). The statistical analysis by One-way ANOVA of Repeated measures depicted highly-significant results for A549 cells at 12 hr (p < 0.0001); 24 hr (p < 0.0001) and for HepG2 cells, at 12 hr (p value of 0.0007). Furthermore, against this setup, we found that 24 hr aseptic-storage of coated-plates enriched the response in cell-culture viz. the morphology and viability in (both) cells, speculating enhanced cell-adherence on stored coated-pRS-1 (Fig. S8C and F). The statistical analysis for A549 cells was found to be significant (p < 0.05). The HepG2 cells showed a significant (p = 0.0097) trend where viability was initially high and restored to a normal-value with the highest conc.-gradient (20.0 µg/ml).

Comparative analysis towards cell-viability revealed a positive but heterogeneous-trendline, distinctly, HepG2 cells depicted an overall >200% viable-cells at 12 hr, 24 hr and 24 hr-precoated-plates (Fig. 3A), Two-way ANOVA in Turkey’s Multiple comparisons for 12 hr depicted significance at 2.5 and 20.0 µg/ml (Fig. 3A) coated-protein (p value = 0.0467) whereby, the A549 cells were found to show efficient growth with 200% increased-viability, especially at 24 hr-precoated-plates (Fig. 3B). The statistical analyses for A549 cell-viability at 12 hr and 24 hr were significant as assessed by Two-way ANOVA (p < 0.0042 to 0.0001) and Turkey’s Multiple comparison test (p < 0.0001). To further understand reasons towards a lowered cell-viability at 24 hrs in the second setup, we studied signs of probable apoptosis or cell-death through Flow cytometry, Annexin V-PI analysis, in A549 cells. A549 cells have a slow growth as against HepG2 cells, thus we wanted to ascertain their response. At 2.5; 20.0 µg/ml coated pRS-1, after > 24 hr A549 cells revealed no cell-death. Also, no-signs of late-apoptosis were observable (results not shown). As per ATCC MTT-cell assay also manifest (https://www.atcc.org/~/media/DA5285A1F52C414E864C966FD78C9A79.ashx) cell proliferation-rate and not just a suspected cell-death. Thus, an assertive cell-viability observed for cells-cultured on pRS-1 coated-plates, stored for ∼24 hours, showing absence of apoptotic cell-death, connotes no cytotoxicity-associated so far, with *in vitro* cell culture. Concluding from these observations, over a period of ∼12 hrs cells show >80% viability and upto ∼24 hrs, cells had the normal cellular-morphology and proliferation-rate.

**Fig. 3.**
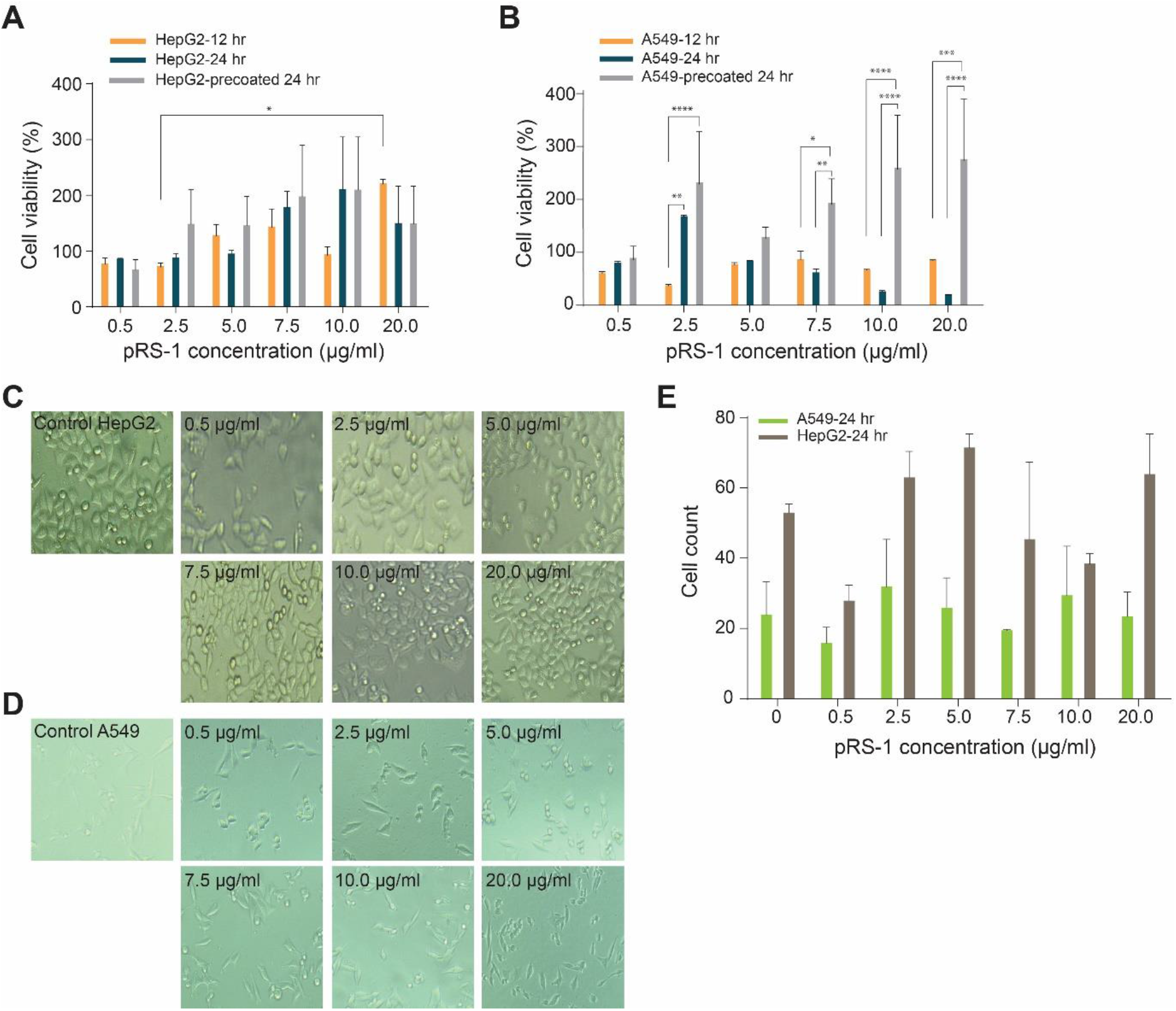
Cell-Viability analysis shown through graphical comparison of three-set of coating studies; 12H, 24H & the 24H precoated plates. (A) HepG2 cells. (B) A549 cells. Cell-adhesion in HepG2 & A549 cells with increasing-gradient of coated pRS-1, cultured upto 24 hrs in pRS-1 precoated plates (Nikon). (C) HepG2 cells and (D) A549 cells. Representative images showing cell densities of pRS-1 treated cells (original magnification 400x). (E) Graphical representation of cell counts as per GraphPad Prism V. 7, (n=3; p < 0.0001; * = 0.01 to 0.05, significant; ** = 0.001 to 0.01 very significant; *** = 0.0001 to 0.001, extremely significant; **** = < 0.0001, extremely significant).

To gain more insight on stored coated-plates, we quantitatively analysed (their effect on) cell-adhesion through cell-counts after 24 hr culture (Fig. 3C). A significant response was observed (p value < 0.0001), although/apparently heterogeneous; HepG2 cells, with smaller generation-time showed heightened-counts with increased-gradient pRS-1 than controls, likewise, counts for slow-growing A549 cells were also found to exceed the controls (Fig. 3D).

### 3.4. In vivo characterization of pRS-1 in full-thickness, cutaneous wound-closure in diabetic rabbit-ear

*Oryctolagus cuniculus,* New Zealand white rabbits (8-10 weeks; 1.5-2.0 kg) experimental diabetic-models were successfully developed using single and multiple-dose alloxan-administration (Supplementary Fig. S9) and confirmed thereafter (Supplementary Fig. S10). Fig. S9B, (see Supplementary) shows representative Blood-glucose profiles with established hyperglycemic-response within first 24 hr of alloxan treatment, consistently. We observed a high-mortality especially in male-models; although clear-hyperglycemia was evidenced after 24 hr in single-dose and upon 48-72 hrs in multiple-dose regime. As an apt wound healing-model, rabbit ear-ulcer model was selected which ensures (the observation of) contraction-free healing, as against rodents and small animals.

Collagen primarily appears into the mammalian-ECM in early-weeks post-wounding and prepares for next necessary modulation. In this context, we tested our hypothesis of wound-healing with pRS-1-augmentation into healing wound-beds about 3-4 days after injury. Fig S9A (Supplementary), shows schematically, major-steps taken to devise full-thickness bed-cartoons sketched on a diabetic-model H&E-section, and Fig. S9C (Supplementary) depicts, 2.5; 6.0 mm wound-models, in representative control and diabetic rabbit-ears.

#### Young-diabetic group cutaneous-wound healing with pRS-1 augmentation

This group structured the healing pattern in 6.0 mm wound-beds in age-matched, treated-groups against untreated-controls each, with healthy- and diabetic-models, besides the Matrigel-treated diabetic-models (positive control), Table S3 (see Supplementary) depicts study-groups. *In vivo* wound-closure analyses breakdown as the gross-analysis followed by image-morphometry and wound-histology (excised-biopsies). Firstly, the Day-wise gross- analysis (Fig. 4A) depict similar wound-edges with initial-contraction in wound-area by day- 3^rd^. On day 5^th^, pRS-1 (treatment) was administered. Wound-area reduction is clearly observed through day-10 in Matrigel-treated beds; the treated- and untreated-diabetic punched-beds are apparently similar; although treated healthy-models healed earlier than untreated(-healthy) controls.

**Fig. 4.**
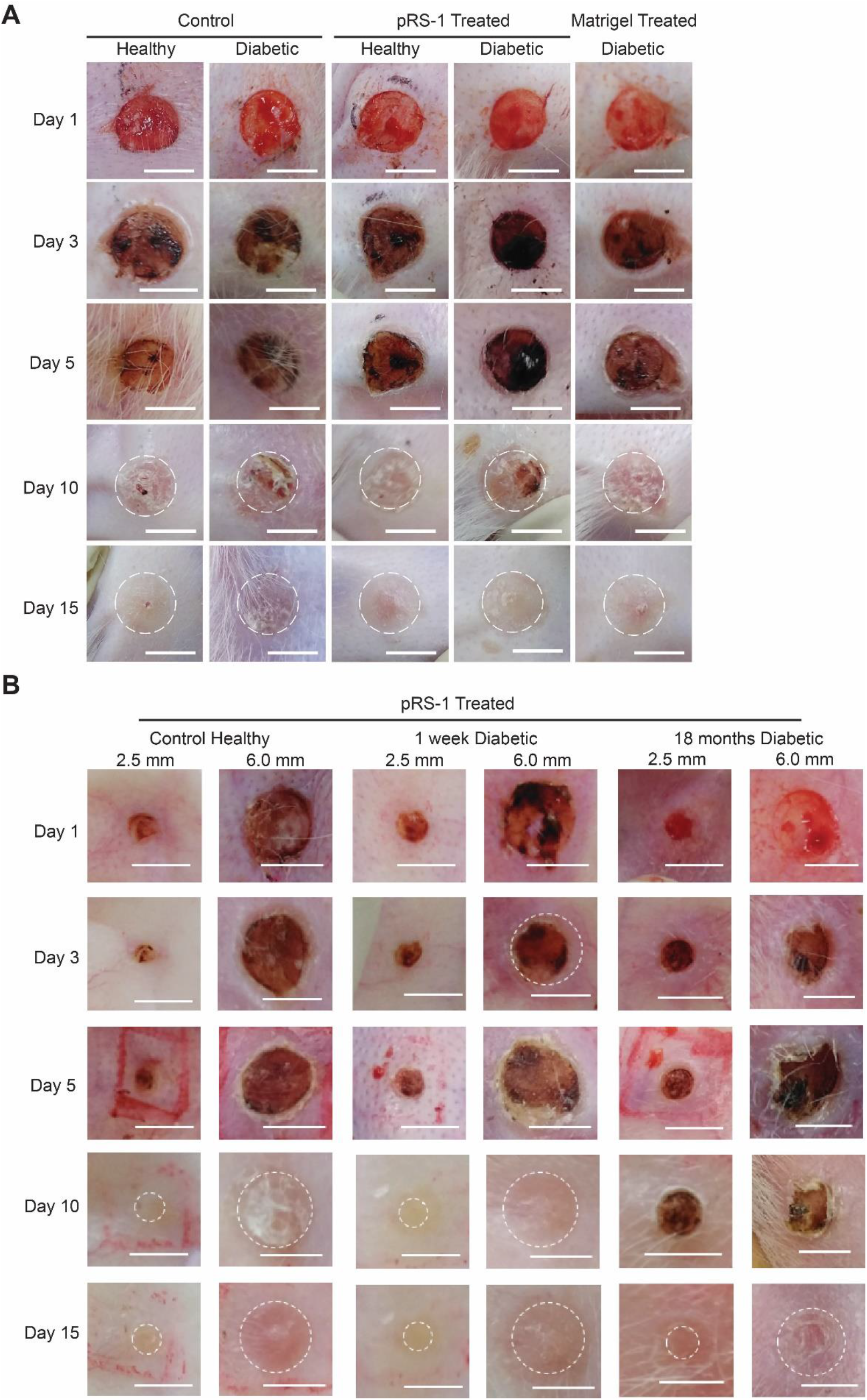
Wound-bed images showing pRS-1 (Recombinant protein) treated wound closure. (A) Young-diabetic group, age-matched healthy and diabetic- untreated-controls; pRS-1 treated and Matrigel augmented positive-controls; 6.0 mm. (B) (pRS-1 treated) Old-diabetic group, assessment between 1 week and 18 months diabetic models.

Secondly, Day-wise image-morphometry was evaluated through a calibration achieved for intensity-change across individual wound-edges (Supplementary Fig. S11). Line-profiles (Fig. 5A(i)) for days 3; 5 depict similar intensity of wound-area, although until day-15, treated-healthy and diabetic-beds show reduced threshold-intensity analogous to Matrigel- group, through the calibration for baseline value (as against untreated-groups). Similarly, the Group-wise line-profiles for treated-groups show closure as Matrigel-group (Fig. 5A(ii)).

**Fig. 5A.**
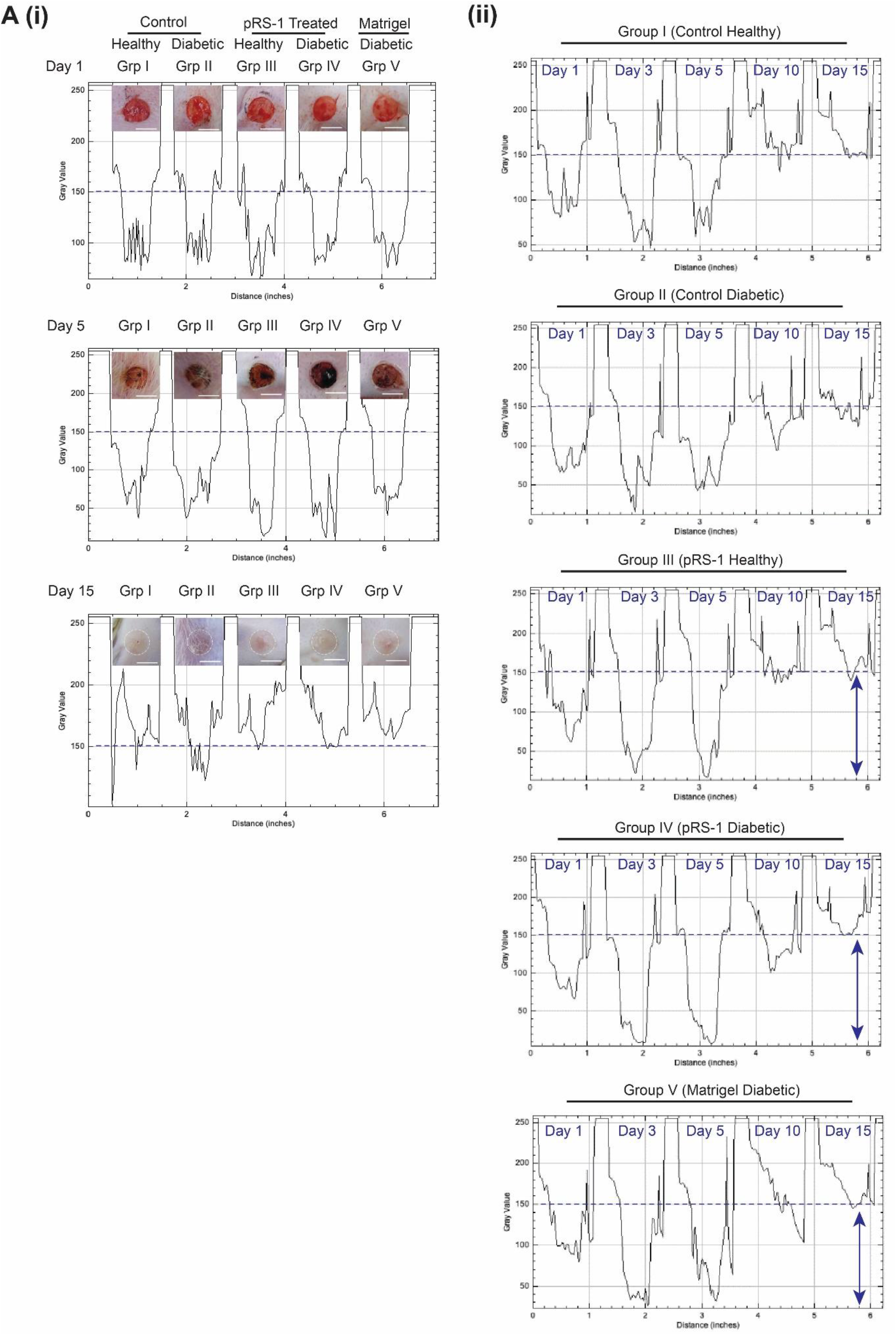
Young-diabetic group wound closure, Blue dotted line depicts the baseline values in Line plots. A(i) Day-wise Line profile: Day 1, 5 & 15 with significant closure in pRS-1 treated Group III & Group IV similar to the value of group V, positive control. A(ii) Group-wise Line profile: Day 15 profile delineates significant reduction in wound-intensity in pRS-1 treated groups, III & IV similar to the positive control, Matrigel treated Group V.

**Fig. 5B.**
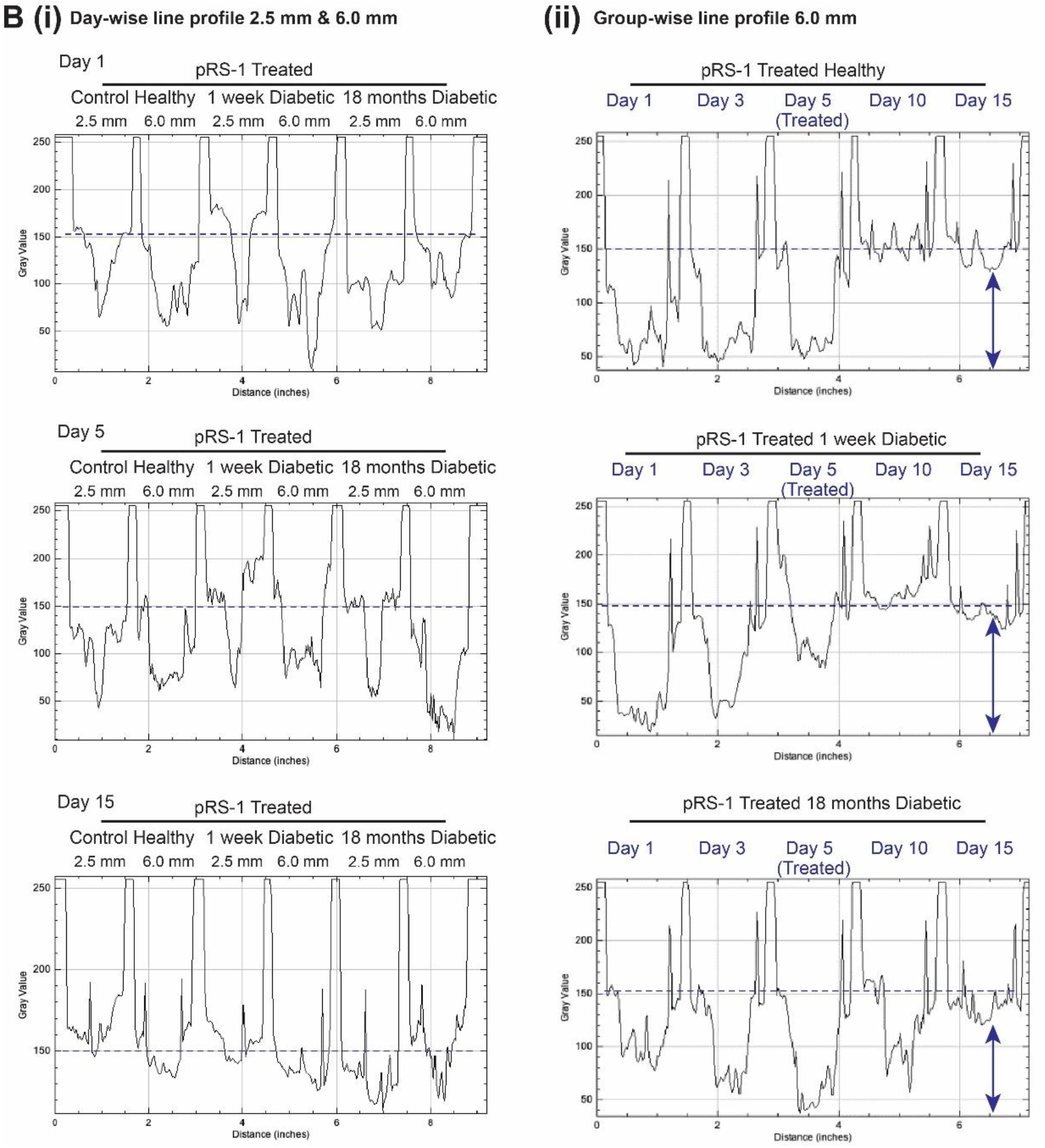
pRS-1 treated Old-diabetic group wound closure, Blue dotted line depicts the baseline values in Line plots. B(i) Day-wise Line profile: Day 15 shows significant closure in 2.5 & 6.0 mm wounds, with slight delay in 18 months group. B(ii) 6.0 mm Group-wise Line profile: Day 15, 18 months-Diabetic profile clearly depicts complete wound closure in line with the 1 week- Diabetic.

Thirdly, histological assessment of excised closed-wounds were evaluated using H&E and differential Gomori’s trichome stains. For reference of wound-closure parameter assessments, the wound-edges were included in excised-tissue using slightly larger punch- tools (8.0 mm). H&E photomicrographs of young-diabetic group significantly show a completely regenerated epithelium/re-epithelialization, without any scab-formation.

Specifically, in comparison to control-untreated the treated-healthy and treated-diabetic sections show very distinct re-epithelialization zones (Fig. 6A, H&E, indicated in yellow arrow-heads), as seen in Matrigel group. Through Gomori’s trichome-sections, just as evidenced for Matrigel-group, the treated-healthy and treated-diabetic groups also depict extracellular-matrix deposition and thick collagen-fiber organization with less fibroblasts in comparison to untreated-controls (Fig. 6A, Gomori’s).

**Fig. 6.**
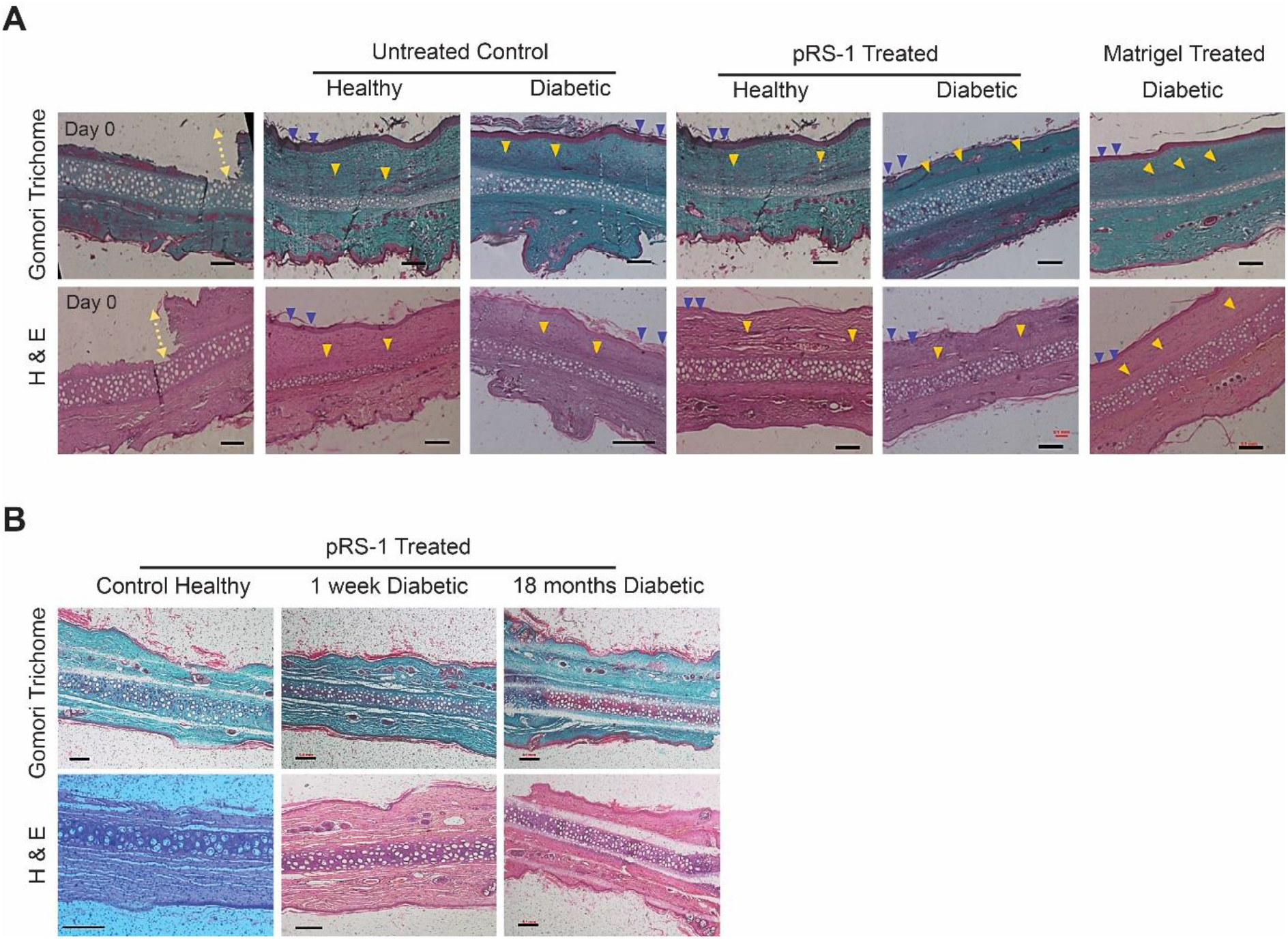
Photomicrographs of Gomori’s trichome and H&E-stained, cutaneous wound-sections of New Zealand White rabbit ear wound-closure biopsies. (A) Young-diabetic group (age-matched) after wound-closure. First panel shows the Day 0 created full-thickness wound-model with intact cartilage- bed; Untreated controls; pRS-1 Treated healthy and diabetic and Matrigel-treated diabetic sections depicting a healthy granulation tissue with re-epithelialization shown in yellow arrow-heads, completely closed epithelium is marked in blue arrow-heads showing complete wound closure and absence of any signs of inflammation, scarred tissue. (B) Old-diabetic group. The healthy, 1 week diabetic tissue-sections reveal apparently similar attributes, as normal granulation tissue without any signs of inflammation as the Young-group although the 18 months diabetic sections showed wound- closure slightly late but pRS-1 effected in an overall normal healed-tissue (n = 3); Scale bar= 0.1 mm.

This age-matched group for wound-closure depicted a similar temporal pattern of healing among the groups (in major observations), thus prompting (us) to study pRS-1 response in a sustained, old-diabetic model (age-differences) as it could elaborate effects of long-standing diabetes in wound healing.

#### Old-diabetic group analysis towards effectual wound-closure

We analysed wound-closure in old-diabetic group, constituted of young ∼1 week diabetic, and 18 months, older diabetic-models and healthy-controls where all groups received pRS-1 treatment. Besides 6.0 mm wound-beds, in this setup 2.5 mm punched-models were also included (Fig. 4B).

After treatment, Day-wise analysis depict significant differences in healing-patterns, specifically in old-diabetic models. 2.5 mm wounds, in young-diabetic (1 week) and control-group, from day 2^nd^-5^th^ are similar (Fig. 4B, Day1-5); and a faster-closure can be seen after pRS-1 administration, until day-9 (in both models). Whereas, in 18 months (long-term) diabetic-2.5 mm models, a delayed-closure with least changes until day-7 (treatment-day) and a sharp decline in wound-area with complete-closure is observed only after day-12 (Fig. 4B, Day 15). Additionally, in young-diabetic, 6.0 mm diabetic-wounds reveal a more-dry, flaky gross-appearance; and gain complete-closure after days 12-15 (Fig. 4B, Day 15) as against control, which recovered by day-9. Whereas, in long-term diabetic-6.0 mm wound, characteristic diabetic-conditions were depicted. Complete-closure was followed as delineated by 2.5 mm models; however, in this group, models achieved closure after overall 15-17 days (Fig. 4B, Day 15).

Differential stain, Picro Sirius red (PS) was employed to characterize diabetic-wound collagen-content and observe both, qualitative and quantitative manifestation of a healthy wound-closure, whereby the appearance of collagen type III-thin fibres (stained yellow-green) along with the type I-thick fibres (stained deep-red) occurs in a healthy granulation tissue. In the young-group, we observed treated group-sections normal tissue-like texture with clear yellow-green stained regions (which were highlighted through the polarised microscopy) as seen in Fig. 7A (yellow arrow heads), blended conspicuously in the whole dermal-tissue with deep-red type I, thick bundles, both in treated healthy- and diabetic-tissues (Group III & IV); whereas, the apparent deep-red fibres exclusively in diabetic-sections (Group II). Notably in these sections, Group III & IV, the healthy tissue can be seen after pRS-1 assisted wound-closure similar in texture to that of the control healthy-section in Group I as shown in the histogram, Fig. 7B and the quantitative-analysis (Red intensity) of the stained-sections Fig. 7C. Sustainably, the stain can be used to characterize the healthy- tissue in examining diabetic-histology accurately and pRS-1 seems to augment and synergise the natural process of healing where red intensity of treated diabetic (Group IV- 139) is lower than the untreated diabetic (Group II-163), expediting the experimental diabetic-healing as seen in the healthy-models without any scar tissue or inflammation as detailed in Fig. S13 and Fig. S14. Further, in Group V, Matrigel sections show thick-fibres exclusively and some thin-fibres, indicative of the diabetic-tissue as the red intensity is higher (169) with low yellow-green type III fibre-content. Thus, the response seen in the pRS-1 treated-models seems to yield to a healthy collagen-ratio in the closed wounds.

**Fig. 7.**
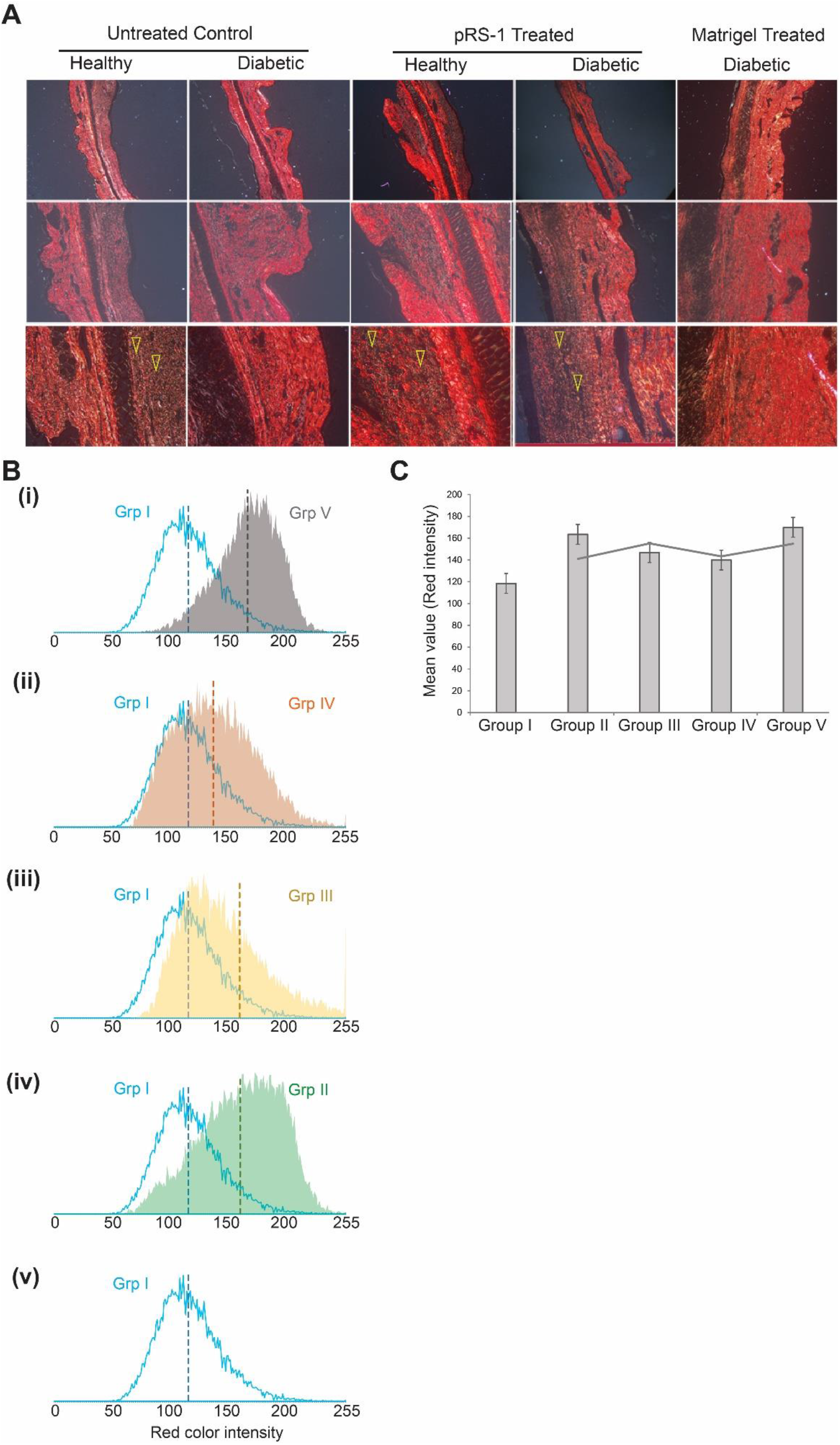
Picro Sirius Red stained-sections of Young-diabetic group. (A) Plane polarized images marked in yellow (arrow-heads) in Control-Healthy, pRS-1 treated Healthy- and Diabetic, cutaneous tissue showing the yellow-green birefringent collagen type III fibres as against the Control-Diabetic and Matrigel Treated Diabetic tissue sections (Original magnification from upper panel-below: 4x, 10x, 20x; Olympus). (B) Quantitation of collagen-content shown as the Red- intensity (mean, ImageJ) plotted in histogram whereby Group I (Control Healthy), Group III & IV (pRS-1 treated-Healthy and -Diabetic) intensity/collagen-content apparently is similar and is less skewed (and indiactes a healthy-texture with mixed yellow-green type III collagen-fibres therefore less red-intensity) as against Group II (Control Diabetic) and Group V (Matrigel treated) which exclusively show deep red type I collagen and absence of yellow-green type III collagen showing more skewed histogram. (C) ImageJ Red Intensity bar graph of the groups indicating treated Groups III (Healthy) and IV (Diabetic) show collagen-content through the red-intensity restored to the levels seen in Group I (Control Healthy).

Further, Day-wise image-morphometry evaluated through Line-profiles (ImageJ) calibration for intensity-change across individual wound-edges show much faster closure in the 2.5 mm than 6.0 mm wounds (Fig. 5B(i)). We studied the Group-wise healing through Line-profiles (Fig. 5B(ii)) and Plot-profiles (ImageJ) for 6.0 mm-punch (Fig. S12).

The Line-profiles (Fig. 5B(ii)) after treatment, depict similar intensity-changes in control- and young-diabetic (1 wk) wound-area, and notably, the 18 months diabetic-wound also exhibit faster healing.

Here, control and young-diabetic Plot-profiles for day-15 (see Supplementary Fig. S12) show evident wound-area reduction and complete-closure as illustrated in the similar wound-bed intensity as surrounding un-wounded tissue, however, 18 months-diabetic day-15 Plot- profile shows almost contracted-wound bed through the contoured area (with high intensity indicating the still-healing wound). Lastly, H&E histo-photomicrographs (Fig. 6B, H&E) of treated young-diabetic and control wounds reveal a similar texture in terms of re-epithelialization in absence of any scar-tissues. The old-diabetic wound shows complete epithelialization by day-15, although epithelial-tongue is irregularly patched with many grooves. Day-15 Gomori’s trichome-section (Fig. 6B, Gomori’s) depict healthy granulation-tissue in control and 1 week diabetic-wounds, whereas in old-diabetic (18 months) the immature granulation-tissue with less organized collagen signifies delayed wound-healing.

### 3.5. In silico modelling of atomistic interaction between Scl1 and human-integrin

We also attempted to understand if Scl1 can interact with the human-integrin protein the same way as human collagen does. We generated complexes of Scl1-peptide stretches with human-integrin (as described in section 2.9.2) and compared the interaction pattern with human collagen-integrin complex reported in PDB. Comparative analyses of intermolecular interactions and binding energies of the CL-domain trimeric-peptides from Scl1 and human collagen with human-integrin revealed very similar pattern (Fig. S15). The key residues of human-integrin involved in human collagen binding were N154, Y157, D160, Q215, D219, S257, H258, Y285 and N287, which mostly formed H-bonds, aromatic H-bonds, salt bridges and π-interactions with human collagen residues (Fig. 8A and Fig. S16). The Scl1-peptides too interacted with all the aforementioned key residues of human-integrin and additionally with I156, E256, G260, S261, V282, R288, D292 and E299. Table 2 lists all the interaction partners from human-integrin and the three monomeric chains of Scl1-peptides and human collagen.

**Table 2.** Residue interactions of 10 different Scl1-peptides and human collagen, with human- integrin protein.

| Residue stretch | Interacting residues of human Integrin | Interacting residues of Scl1 residue stretches |  |  |
| --- | --- | --- | --- | --- |
|  | Chain A | Chain B | Chain C | Chain D |
| S1 | N154, S155, Y157, D160, D219, V282, N287, R288 | K74, G75, D76, R77, E79 | G78, E79, T80 | S71 |
| S2 | N154, I156, Y157, D160, D219, E256, S261, L286, E299 | R95, G96, Q99, K101 | Q98, K101, R104 | E106 |
| S3 | N154, S155, Y157, D160, Q215, D219, H258, S261, N287 | E119, R120, E122, K126 | E119, R120, E122, K126 | E128 |
| S4 | N154, Y157, E256, S257, H258, Y285, N287 | A137, K139, Q143 | Q140, Q143, A136 | - |
| S5 | N154, Y157, D219, E256, S257, H258, S261, D292 | P158, K161, K164, R167 | K161, K164, G165, R167 | E169 |
| S6 | N154, Y157, D219, S257, H258, S261, N287, R288 | Q179, K182, E184 | K182, E184, T185, Q188 | Q179 |
| S7 | N154, S155, Y157, Q215, D219, S257, H258, S261, R288 | P200, Q203 | E202, Q203, A206, G207, Q209 | E211 |
| S8 | N154, Y157, S257, H258, G260, S261, Y285, N287, R288 | P220, E221, S223, P224, G226 | P224, E227, Q230 | E225, G232 |
| S9 | N154, Y157, D160, D219, S257, H258, G260, S261, N287, R288 | V242, T243, A245 | A245, E247, K248, D251 | S239, E253 |
| S10 | N154, Y157, D160, E256, H258, S261, N287, D292 | G263, A266, V270 | A266, T268, P269, N272 | K267, H274 |
| Human collagen | N154, Y157, D160, Q215, D219, S257, H258, Y285, N287 | HP6, F8, E11, R12 | F8, HP9, E11, HP15 | - |

**Fig. 8.**
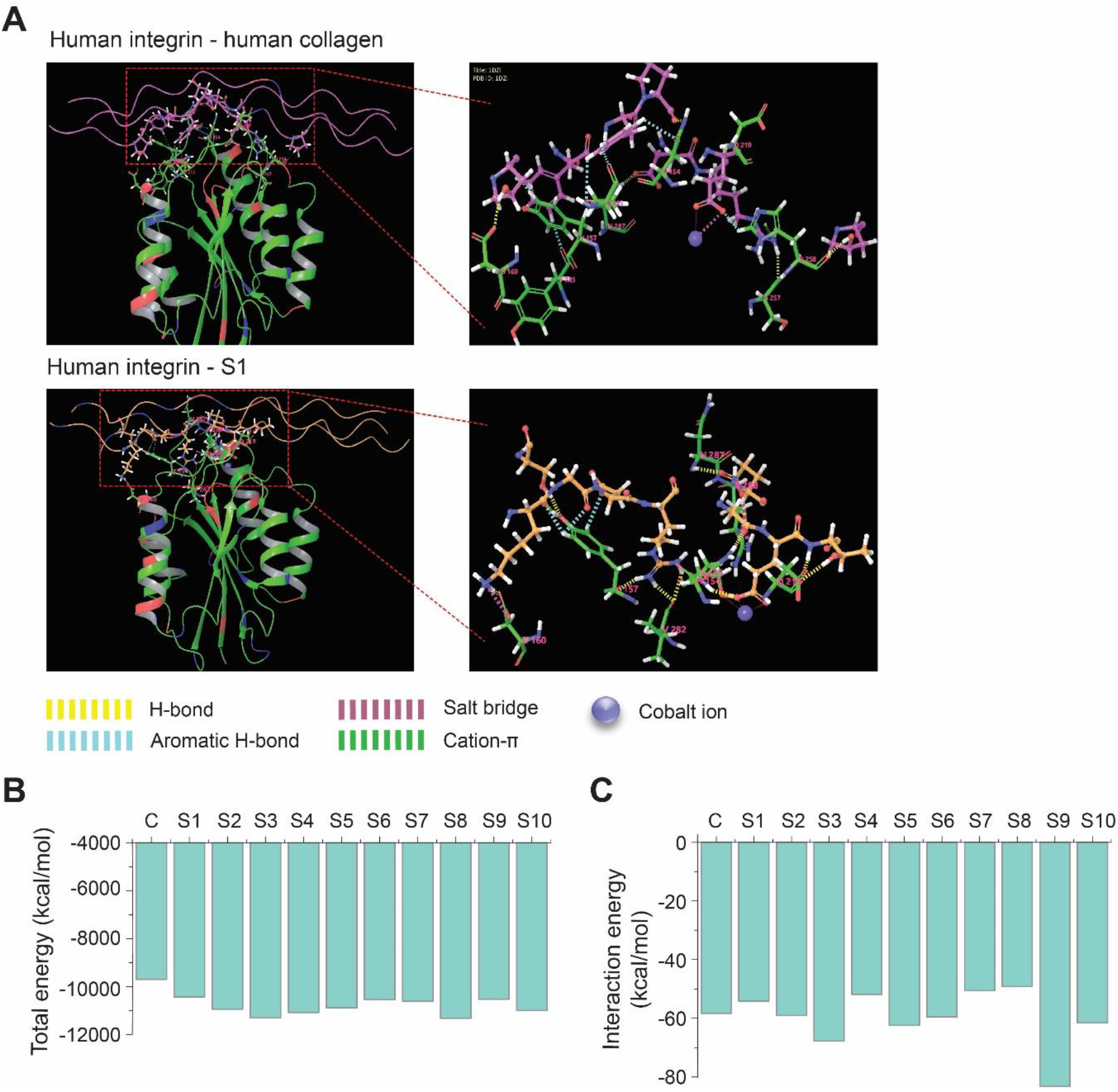
*In silico* atomistic modelling of Scl1-peptides and their interactions with human integrin receptor. (A) Non-covalent interactions of human integrin (green) with human collagen (pink) and a peptide S1 from the Scl1 collagen-like domain (orange), (B) Total- and interaction-energies of the human integrin-human collagen (Denoted in the graph X-axis by C) and human integrin-Scl1 peptide complexes (Denoted in the graph X-axis by S1-S10).

The total energies of all the human integrin-Scl1 peptides were observed to be slightly lower (ranging from -10428.57 to -11320.74 kcal/mol) as compared to the human integrin- human collagen peptide (−9699.32 kcal/mol), of same length. The overall stabilization energy (interaction energy) between human-integrin and human collagen as calculated from the reported structure (pdb id: 1DZI) was -58.32 kcal/mol, while the stabilization energies of the Scl1-peptides ranged from -49.22 to -83.26 (average value from 10 complexes: -59.94). The trimeric-peptides S2, S3, S5, S6, S9 and S10 showed better binding than collagen. The total energies of the complexes and interaction energies of collagen and Scl1-peptide (S1-S10) with human-integrin have been plotted in Fig. 8B and C. The comparative analyses indicate better interactions and enthalpic stabilities of the human integrin-Scl1 peptide complexes as compared to the human integrin-collagen complex.

With the sub-cutaneous supplementation of pRS-1 which is a collagen-mimic we have not observed any untoward effects in the tissue histology. Notably, there is complete absence of any hypertrophic scarring that could result with excessive collagen in the healing-tissues and thus indicates the safe application of this recombinant protein in the animal-wound models.

## 4. Discussion

Streptococcus pyogenes or GAS (Group A Streptococcus), a human specific pathogen has imminent concerns in terms of a high disease burden. The enhanced pathogenic-potential of GAS is mediated through the ensemble of many virulent-markers, most important are adhesins which define primarily fate of the assault as mediators in initial stages of establishment of infection. Recently, the streptococcal collagen-like proteins (Scls) have given insights toward novel potential pathogenic-adhesins of GAS mediating internalization through matrix interaction [38],[39], associated human tissue-protein cryptic-interactions, i.e., collagen types I and IV cross-reactivity, and a sero-positive [11], among its important pathogenic facets.

In the similar context, emphasizing their essential roles in GAS pathogenesis, in our previous studies, we found conservation of Scls among 27 *emm* types identified in clinical- GAS isolates [11]. Many microbial species carry collagen-like (CL) sequences in their genome, among these, streptococcal-proteins received more evaluation in the last decade.

As an evolutionarily conserved protein domain, the collagenous-domain is evidently expressed by many distant life-forms, it’s triple helical structure lends (it) the name ‘trimer’[4],[5]. In this context evaluating CLs, recently, certain biophysical studies of Scls have shown a glimpse into their structure, stability and variations to their corresponding human-proteins. Yu et al., depicted role of charge-distribution, preference to non-proline residues in a fraction of CL-domains in Scl2, evolutionarily potentiating folding and imitating human-trimer structural feature [3]. Some studies have listed the essential inherent features in Scls akin to human system of collagen(s) [1],[41] as their pH and thermal-transitions reflect profiles as mammalian triple-helices; and these depict the bacterial CL-domain phenotype as the major human fibrillar-collagen, type I [1]. Distinctly, the studies have listed recombinant Scl2-constructs from varied GAS strains and characterized their stability from biophysical point of view [42]. The discreet structural constraint on bacterial proteins to achieve closely- bound, yet classical-(trimer) repeats found in (fibrillar-human) collagen hints at its specific similarity in structural-feature, and expression of integrin-binding site as the functional- insignia to collagen. The diversified, evolutionarily-conserved mammalian collagen are not alone in nature, markedly the microbial-proteins have been associated with bearing similar sequences as collagen-mimics.

In spite of close association to mammalian protein features, the bacterial-protein lacks post-translationally modified-residues due to it’s evolutionary-biochemical confines. These/[The (bacterial) CL-proteins] are stable through varied electrostatic-interactions; polar residues; higher imino acid, proline content; hydration-networks and standard-helical V- domain with critical implications towards folding (of CL-proteins) [1],[43]. A very recent study by Lukomski group indicated Fn-interaction of GAS Scl1 and not Scl2-V-domain models through complementation-studies relating the V-domain pathogenic potential, which was previously unknown [44]. These features have a benefit aptly rendered to the molecular- versions of CL-proteins, expressed in bacterial systems, which are stabilized through certain mechanisms other than known in human collagen [42],[41],[1]. Reviewing on these excerpts we selected GAS serotypes with varied virulence for this study.

Towards sequence characterization of Scl(s) we included, GAS M1 [23, 45], extensively studied rheumatogenic-serotype; GAS M1-2 an M1-variant, isolated from our previous studies [12, 46]; and nephritogenic-serotype M49 [47], also the smallest of GAS sequenced- genomes. In serotypes with distinct-virulence, expressed-adhesins Scl1, Scl2 reflect semblance in domain-structures and we found assertive homology with variations in certain primary aa sequence and (in) number of CL-repeats, as also reported in earlier characterizations [23]. This essentially corroborates our previous observations where we deciphered ubiquity of Scls among GAS serotypes [11]. Multiple sequence alignment of the GAS M1 and M49 Scl1 proteins show 77% sequence identity, but M49-Scl1 has a 63-residue insertion at the end of the CL-domain, which is not present in M1-Scl1. This insertion is a triple repeat of a mostly negatively-charged “EKSPEVTPTPETPEQPGEKAP” stretch. Similarly, global alignment of GAS M1 and M49 Scl2 proteins reveal 71% sequence identity; however, a 42-residue insertion rich in negatively-charged residues (D and E), was found at the beginning of the CL-domain of M49 Scl2. Also, another 18-residue insertion with a repetitive GxD (GKD or GQD) triplets was found in M49 Scl2 towards the end of the CL- domain. Such insertions might be specific for a certain strain and indicate the strain specific CL-domain features in GAS; although, CL-domain length and primary sequence is strain- dependent but invariably, collagen-like domain stability is unaffected by variation in their residues [42].

Henceforth, M1 Scl1-protein was selected for characterization. Full-length construct (813 bp) with N-terminal V-domain which has role in CL-polymerization [48], [43],[49], followed by the CL- and L-domains, yields a soluble protein pRS-1 (rScl1) in the strong pET- expression system. We focused to analyse the relevance of full-length collagen mimetic- identity of Scl1 to human ECM-protein as a prelude to its bio-activity, thus V-CL (339 bp) M1 GAS *scl1* smaller-construct (in pQE-system) was not further included in this work. Although studies have reported Scl2 recombinant-protein constructs [42],[43] like V-domain with tandem-repeats of bacterial-CL to achieve a large fibrillar molecule (as collagen); yet a full-length construct towards characterization of its effects *in vivo* has not been shown.

Apparently, the observed mass and migration behaviour on denatured-gel and /or purification of bacterial-surface proteins have certain anomaly, in terms of difference in observed mass and calculated value, analogous reports have been presented for recombinant Scl-constructs [40],[50]. Similarly, pRS-1 migrated on a heavier-side on SDS-gel than its predicted mass (from protein-sequence analysis). Towards initial characterization, *in silico* features of the construct through Protparam tool (Expasy), showed it stable with its major domains. While optimizing protein quantitation, we monitored rScl1 for its residue- proportions which revealed less conjugate-amino acid residues thereby making the estimation-procedures ineffective.

Consistent with the observations that Scl(s) are mimics of human collagen [41], homology analysis through BLAST tool reflected a 50-55% identity of collagen-like (CL-) domain of Scl1 (cds) to human-collagen type I α-chain (position 162-1218, propeptide domains-free). In our *in silico* structural rScl1 model, total energy of the modelled monomeric-unit was calculated to be -7359 kcal/mol while that of the trimeric-assembly was -23549.6 kcal/mol. This quantitatively showed that the collagen-like trimer form is thermodynamically more stable than the monomeric form, as sum of total energies of 3 individual monomeric chains (3*(−7359)= −22078.2 kcal/mol) is higher than the total energy of the trimer (−23549.6 kcal/mol). The trimeric-assembly is stabilized by H-bond and salt bridge interaction networks formed between the monomers. Also, polar side chains of the X and Y residues in GXY motifs of one monomeric chain makes H-bond/salt bridge interactions with the main chain of the adjacent chains. Most elusively, the absence of the hydroxyproline residues that are a characteristic predominant sequence-unit in human collagen structure, don’t seem a determinant in structural homology analysis. Convincingly, these initial-analyses hinted at a probabilistic biomaterial aspect of rScl1.

We attempted to correlate two estimation methods; first being Protparam [Expasy] analysis of amino acid-residues; second, the soluble-protein estimation akin to linear- approximation of protein through Coomassie-blue, silver-stain or immunoblot of pRS-1. As previously mentioned, our construct revealed meagre- (0.7% tryptophan-, 1.0% tyrosine-, none cysteine-residues and 1.4% phenylalanine- and 4.0% threonine-)/conjugate-residues, as expected direct-quantitation (spectrophotometric absorbance, molar extinction coefficient of pRS-1 construct)/depicted ambiguous yields. Relative to the peptide-bond absorption, tryptophan is followed by other (phenylalanine, tyrosine and histidine 6x-molar extinction coefficients) amino acids towards their contribution in absorption, which is least in proline; although in a peptide these residues manifest the effects of buffer—the protonation of the carboxylic acid etc. and other conformational effects—as in a peptide, the proline absorption is changed (3x coefficient; except if it’s at the N-terminus) [51, 52]. Importantly, insights on effects of these residues in different buffer systems might further lead to ambiguity. And, we were prompted to select a more robust method and in dearth of basic-residues, pRS-1 construct was undetected by the Bradford-reagent. Among the colourimetric assays, BCA method is the sensitive copper-based assay [53],[54] whereby the amount of reduced Cuprous-ion is indicative of the total protein. It is a more consistent estimation method which yielded stable protein quantitation as it relies on the peptide-bond [55, 56] as against the ambiguous quantitation caused due to absence of certain residues among other methods i.e. Bradford and Lowry assays.

To evaluate our full-length construct features toward distinct roles in *in vitro* and *in vivo* systems we optimized our study in physiological buffers. Notably, the *in vitro* pRS-1 characterization depicted stability and integrity in, Bicarbonate carbonate (B/C, pH 9.6; 0.1 and 0.3 M), Phosphate buffered saline (PBS, pH 7.4; 0.01 M) and Tris (pH 8.0; 0.1M)-buffer, in the order mentioned.

Further, skeptical observations over loss of protein; imminent and more explicit loss of integrity over storage was noted throughout the initial phase of study, in spite of effective requisite measures being observed. Conspicuously, we observed certain aspects similar to a study that quotes storage in PBS revealed serine-mediated aggregation manifesting loss of proteins; and the behaviour observed irrespective of pI of protein and ionic-strength of buffer [57, 58]. Moreover, the stabilizing influence of imidazole was seen in pRS-1-purified, concentrated protein, while buffers without-imidazole (buffer-exchanged) show loss of protein, those with-imidazole had soluble-protein stable on storage for >1 week. Thus, we found the pRS-1 protein more stabilized in B/C buffer than PBS and Tris.

Although, buffer-exchanged, concentrated and filter sterile (0.22 micron) pRS-1 was stable few days over storage; however, we analysed the protein through estimation-assays and SDS-PAGE; correlated them, at most to essentially calibrate a relation of the stored protein concentration with time, and finally, we corroborated both the methods (data not shown). It might be noted that loss through filter-sterilization is a cue as observed in previous studies delineating serine-mediated protein degradation in PBS. To further prep pRS-1- integrity and characterize the CL-repeats we stipulated effect of trypsin and found our construct resistant. It is mentioned previously whereby these domains were inert and the chimeric foreign-residues were found to be sensitive [50]. Notably, delineating enzyme susceptibility, we have found pRS-1 bio-available and amenable to bio-augmentation.

For our study, we wanted to focus on characterization of matrix-like features and non-toxicity, imminent for *in vivo* studies. We studied the specific effects of pRS-1-coating on cells grossly; whereby upto 20 µg/ml pRS-1, proved innocuous; it manifested stimulation in cell growth with increased-concentrations, accompanied by spur in their microscopic appearance. The effects on fast-growing HepG2 and slow-growing A549 cells wholly are heterogenous, although in the three setups; in 12 hr, 24 hr and the third, aseptically coated- stored plates for 24 hrs, we found stored-plates support best results with maximal cell- viabilities. Apparently, collagen-like proteins with the essential functional cues of mammalian collagen as matrix-adherence motifs might guide the bioavailability of pRS-1 through integrin-binding and might synergize the course of natural cell-adherence.

Continuing with this hypothesis verification, we found the stabilizing and direct effects on cell-adhesion through an increased cell-count temporally, parallel with the increased pRS-1 gradients. The concentrations were arbitrarily selected for the assays, although, commercial sources for coated collagen-plates were simulated for reference which depict 10 µg/ml coated-collagen (Collagen-coated culture systems, Sigma). Here it would be notable to elaborate that previously in solid-state binding assays using coated modified-Scl2 fragment, intricate Fn-binding site was reported [50]. However, we elucidated the un-modified rScl1 (pRS-1) bio-active and further—the extensive possibility of stored coated-pRS-1 plates is found effective towards ‘matrix-supplemental’ use. These studies are insightful, although other studies with modified-CL fragments which augment cell-interaction are available [44, 50]; however, the complete full-length construct has not been evaluated towards biomaterial characterization in an animal model.

As Lemo et al. have listed features of the rabbit ear-model, especially the contraction-free healing modelled [59] in it as seen in human wounds, making it an ideal system to study the cutaneous-response [60]; thus, we elucidated our study in the ear-full thickness model in diabetic rabbits. We targeted for stabilized experimental-diabetes in our models, so we maintained them for at least 1 week before initiating the wound-studies. Essentially, single and multiple dosing-regime of alloxan monohydrate successfully resulted in crisp hyperglycemic-models, as referenced [27, 61]; moreover, susceptibility to its diabetogenic but safe-dose has been our marked observation and through careful analysis of literature and experimentally with this study we found the 100 mg/kg intravenous dose safe with less-mortality within the initial week of dosing. This delineates lesser acute damage of this experimental-diabetogenic dose, as was determined from the histologically examined internal organs of these animals; contrasting the higher animal loss within a week with slightly higher doses but inconclusive effects of alloxan detectable in the tissues.

Experimental-diabetes elucidated the essential features of insulin-dependent diabetes mellitus [62, 63],[64], most importantly, altered glycemic-homeostasis; all our models evidenced hyperglycemia effectuating in physiologically gross-increased food and water intake; apparently they were active over the period of experiment. Among the closely followed study strategies, especially factors affecting the generation and maintenance of diabetic-model were closely monitored. Notably, the association of stress-induced hyperglycemic-shock [61] was avoided as dosing was spread over two-days, each animal was acclimated while taking out of cage and until after completed dosing. Apparently, higher mortality ratio observed in rabbits with alloxan was mitigated through a post-administration close, hourly monitoring for blood-glucose and resorting to subcutaneous dextrose-therapy every 2 hrs. Female-models were among the higher-survivors in our groups of age-matched alloxan-experimental diabetics.

The *in vivo* studies were conducted to keep a check at confounding factors that might be introduced through the dosing and subsequent effects through aging, although in our young- diabetic group, affirmative wound closure was observed with similar temporal and histological response of young-diabetic models as seen in the healthy, age-matched control rabbits. Besides, studies from the referenced work, all have a shorter age-span with confirmed hyperglycemic-models; therefore, for our intervention study we deliberated to employ an older diabetic-group as well. Through these evaluations its notable that previous studies; especially, in diabetic rabbit-model, have focused on immediate experimental-hyperglycemia that is achieved within 24 hours (in case of alloxan), as their model of diabetes. It lends major confines [65] and does not delineate the long-term human disease pathophysiology towards wound-closure.

In this context and with the prelude on our young-group response, we formulated the older-group besides young-group, for further study. Among few studies with experimental animal-model of diabetes in rabbit, one group modelled the approach to maintain rabbits with insulin-dosing over a long-term [65], however in our study the old-group animals were sustained for 18 months as per the requirement of study and they manifested the signs of later-stage disease. The physiological parameters of the animals were thoroughly monitored and they were kept clean and dry throughout the study to help protect against any infection, especially through wound-healing phase.

We attempted to evaluate an innocuous synergism in recombinant bacterial collagen-like protein, pRS-1, with natural-physiology of collagen-laying down in healing granulation- tissue. Through our results in the pRS-1 treated young-diabetic model, we have established our hypothesis of administration of recombinant-protein within 3-5 days of wounding to align with natural-collagen laying response (in skin). Further, we were tempted to elucidate the similar hypothesis in older-diabetic (experimental) models as the laying-down of collagen constitutes the primary quotient of a healthy cutaneous-healing. Some studies have reported the significant role of applying their formulation in suspension as the saline. They found that after few hours of local-administration, the upper-layer depicts adsorption preferentially, and saline as against a gel-formulation revealed better absorption/availability in the absorption- kinetics [66]. In this work, we also resorted to supplement pRS-1 as a suspension in PBS and we monitored its safety experimentally and found this suited physiologically.

In initial characterization of our models (old-diabetic models,) we quantitated cutaneous collagen-content using Picrosirius red and to our surprise, it highlighted coarsely bright-red stained type I fibres in 18 months-diabetic full-thickness tissue-sections, as against birefringent green, thin type III collagen in healthy-skin. Notably, this indicates a stark variation of collagen-fibre organization and their proportions in 18 months diabetic(-skin) with thick-fibres exclusively and less-remodelling in contrast to healthy skin. This must infer a more amenable cutaneous-response in the healthy-skin (tissue) reflected through an equal proportion of collagen, indicative of a well-organized/orchestrated collagen-physiology of the remodelled wound, which in hyperglycemia, is disturbed.

Also, distinct thick/flaccid and waxy changes were notable in >1 year-diabetic (ear) skin. Similar observations were stated showing association of scleroderma-like changes with diabetes (presentation) as thickened-waxy skin; (strengthened) distinct-ECM due to glycation, and increased fibroblast synthesis of collagen [67, 68], which are predominately also defined as an insidious and gradual implication in diabetes type I as Scleredema diabeticorum [69].

After pRS-1 treatment in the intervention, the gross wound closure (6.0 mm) was seen in 10-12 days for the young-diabetic group as in the age matched healthy controls; however, the older-18 months-diabetic, healing was delayed. Markedly, in age-matched animals, pRS-1 treated- and Matrigel-groups evidently show a similar gross-healing time and morphometric assessments for wound-closure, and the histology of healed full-thickness cutaneous-tissue concludes complete epithelialization without scar-tissue, no inflammation, and a healthy granulation tissue. We concluded the effects of pRS-1 on similar lines as Matrigel. Thus, with this positive inference towards gross-intervention of pRS-1-treatment in young-models we elaborated to manifest the (gross and tissue-) implications in older-diabetic wounds and additionally we devised, a 2.5 mm wound as (a) temporal control of wound-contraction along with 6.0 mm punch. The older-wounds developed distinct diabetic complications although we detected the observable similarity (of closure) in smaller 2.5 mm cutaneous-beds in both pRS-1 treated 1-week young-diabetic and 18 months older-diabetic models. And, the pRS-1 treated 6.0 mm old-diabetic wound-closure was also found to be augmented as complicated tissue was seen to recover post the treatment-day (and closure was faster). Histologically, despite the conspicuous (effects of) experimental-diabetes in our older-group models, the pRS-1 treated acute wound-tissues shows complete re-epithelialization, distinct new- epidermis with healthy vascularization and hair follicles seen grossly in closed cutaneous- integuments concluding in a normal granulation.

## 5. Conclusion

Collagen-like proteins have been reported from many life-forms with distinct similarities to the conserved nature of the trimer. This insightfully invited many important revelations recently; Scl(s), the pathogenic-adhesins (Scl1 only reported) expressed in GAS connote collagen-structural stability, functional ECM-domains. We have extensively affirmed these similarities *in silico* through stable binding-energy of rScl1-peptides to integrin receptor domains; and experimentally, imploring on recombinant-Scl1 matrix-like features in cell-culture and full-thickness wound as being a safe, non-cytotoxic bio-active collagen- synergist/alternate. Conclusively, diabetic wound-implications in unprecedented > 18 month-old rabbit-model were made; in dearth of studies in models with long-standing diabetes. Examining collagen-quantitation in closed-wound (section/histology), pRS-1 augmented cutaneous-wound has shown a healthy granulation and epithelialization indicating the collagen-ratios, especially in the diabetic-models similar to the healthy-control.

## Declaration of competing interests

There is no conflict to be reported in context to this work.

## Acknowledgements

Indian Council for Medical Research, ICMR is indebtedly credited for the research grant of Senior Research fellowship (SRF) for this work. Dr. Pradip K. Chakraborti, previously affiliated with the Council of Scientific & Industrial Research, Institute of Microbial Technology (CSIR-IMTECH), Chandigarh for their kind gift, pQE30 vector. Mrs. Meenakshi Kaundal from Dept. of Experimental Medicine & Biotechnology, PGIMER, Chandigarh and Miss. Tripti Sahoo from Small Animal House, PGIMER, Chandigarh for their earnest zeal, endurance with the rabbit-models and also their love for the animals; Dr. Vivek Sagar, Community Medicine and School of Public Health, PGIMER, Chandigarh for guidance on cloning, screening; Subendu Sarkar previously affiliated with PGIMER, Chandigarh for initial conception and discussions of Methodology.

## Notes

### Competing Interest Statement

The authors have declared no competing interest.

