## Supplementary figures and Tables for "Synergistic collagen-condiment: Streptococcal collagen-like (Scl) protein in cell-adhesion and diabetic wound-closure matrix": Supplementary File_R Singh etal_BioAxiv.pdf

**Supplementary Table S1. Primer details for the *scl1* gene sequences studied**

| Primer Details | Primer Sequence | Product Size (bp) |
|---|---|---|
| *scl1* | **F:** 5'- GC**G GATCC**gaggtttcttctacgactatga -3'<br>**R:** 5'- GC**GTCGAC**acgtctgtggttgttggcta -3' | 813 |
| *scl1*-CL | **F:** 5'- GC**GGATCC**tattttaaagaagaagattttcaaaaggagct -3'<br>**R:** 5'- GC**GTCGAC**atctttacctggagcgccatctttac -3' | 339 |

**Supplementary Table S2. Animal dose optimizations: Alloxan and Dextrose**

| **Properties** | **Observations & Features** |
|---|---|
| Alloxan Monohydrate Dose | Single dose 150 mg/kg<br>Single dose 100 mg/kg |
| Administration Route | I.V. in saline 4% solution, slow inj. over a minute (on ice) |
| Subcutaneous Dextrose | 5% sterile inj., 10 ml-every 2 h, for ~24 h after Alloxan administration |
| Oral Glucose | Feeding upto 24 h after Alloxan administration |

**Supplementary Table S3. Animal Usage**

| Animal Groups | | | | |
|---|---|---|---|---|
| Healthy Controls | Disease or Negative Control | Wound Healing Study Groups | | |
| | | I (Healthy control, pRS-1 Treated) | II (Diabetic-Model, pRS-1 Treated) | III (Diabetic-model treated with commercial ECM) |
| Male & Female | Male & Female | Male & Female | Male & Female | Male & Female |
| 3 | 3 | 3 | 3 | 3 |

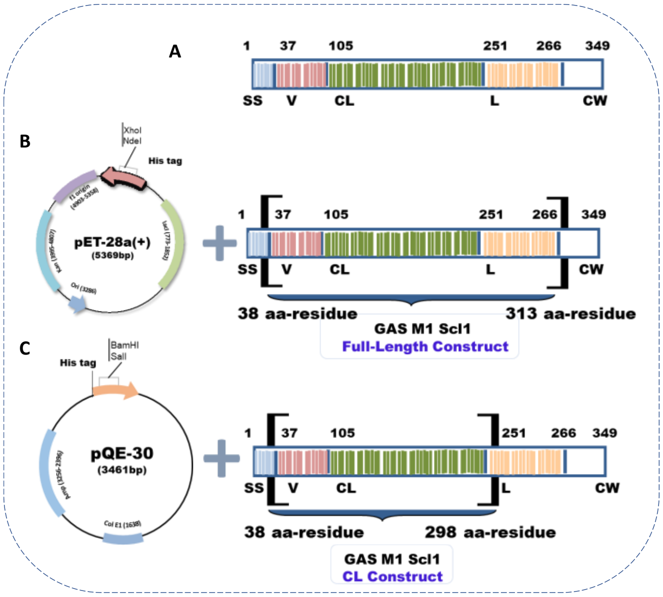


**Fig. S1.** Schematic depicting GAS M1 Scl1 recombinant-constructs, showing major cloned Domains, **(A)** Scl1 complete-domains, signal sequence (SS), Variable-domain (V), Collagen-like domain (CL), Linker (L), and cell wall (CW) anchor conserved in the Gram-positive surface protein cds. **(B)** pRS-1 (recombinant Scl1, 813 bp) and **(C)** pRS-CL (recombinant collagen-like region, 339 bp).

A

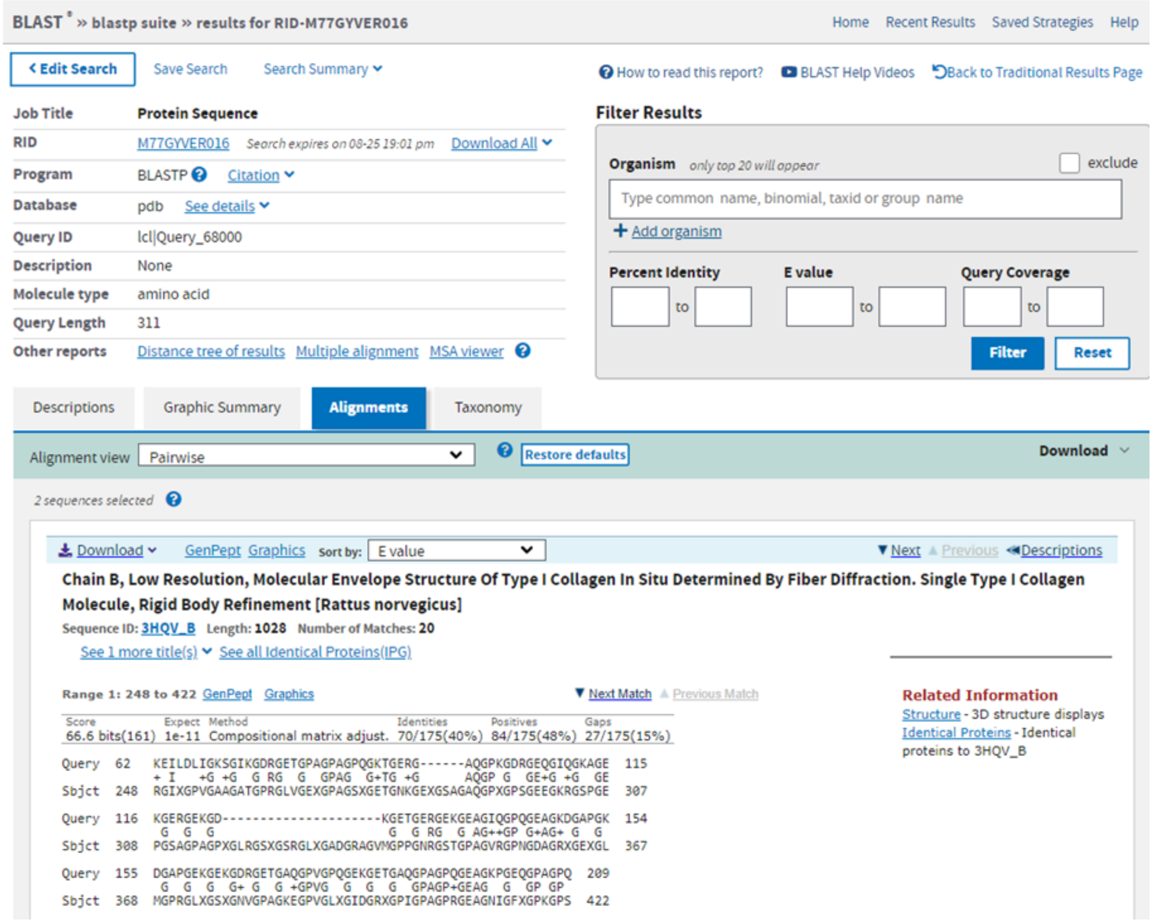


**Fig. S2. (A)** Local sequence alignment of scl1 and 3HQV using BLAST.

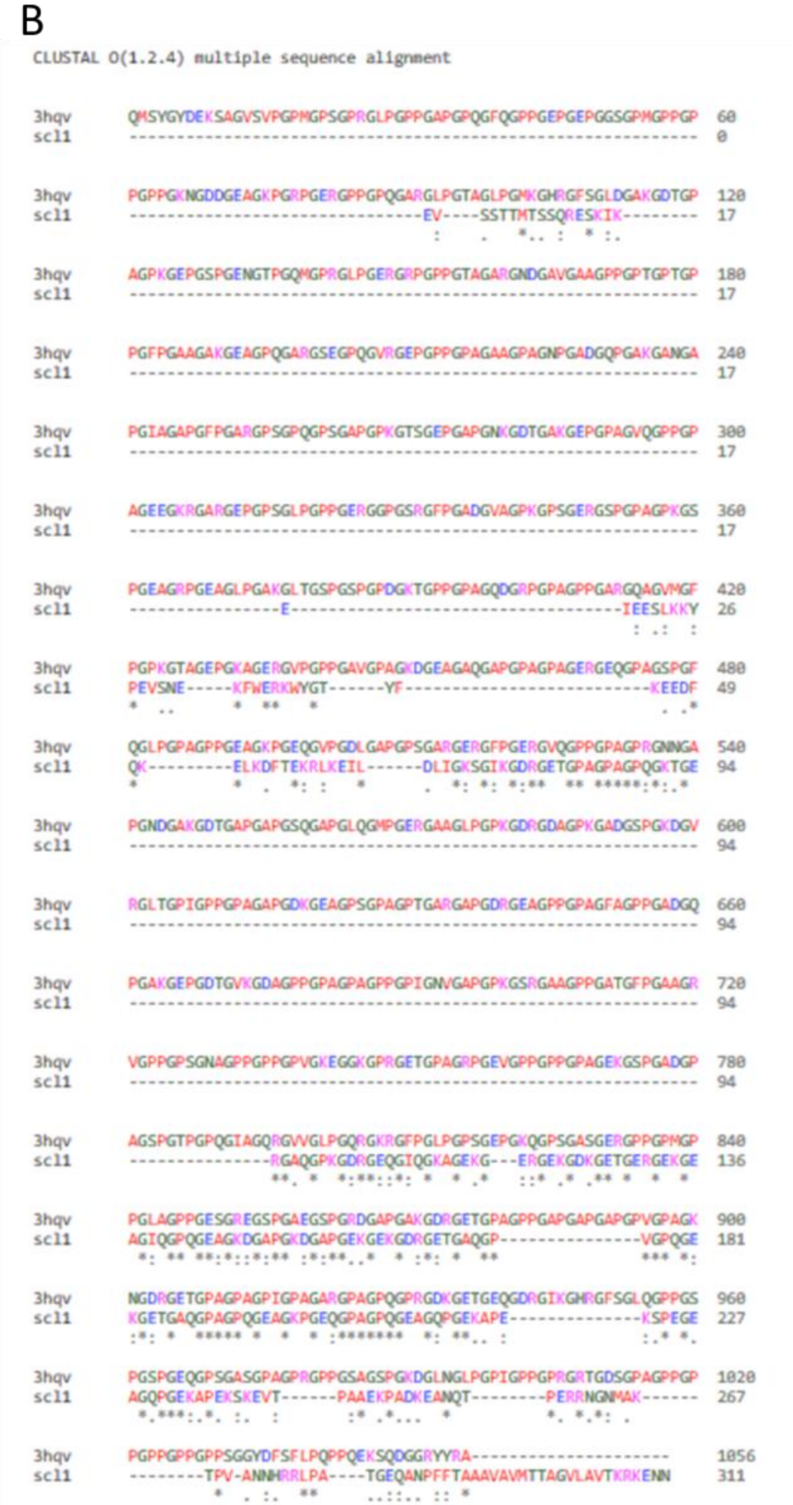


**Fig. S2. (B)** Global sequence alignment of Scl1 and 3HQV with Clustal omega.

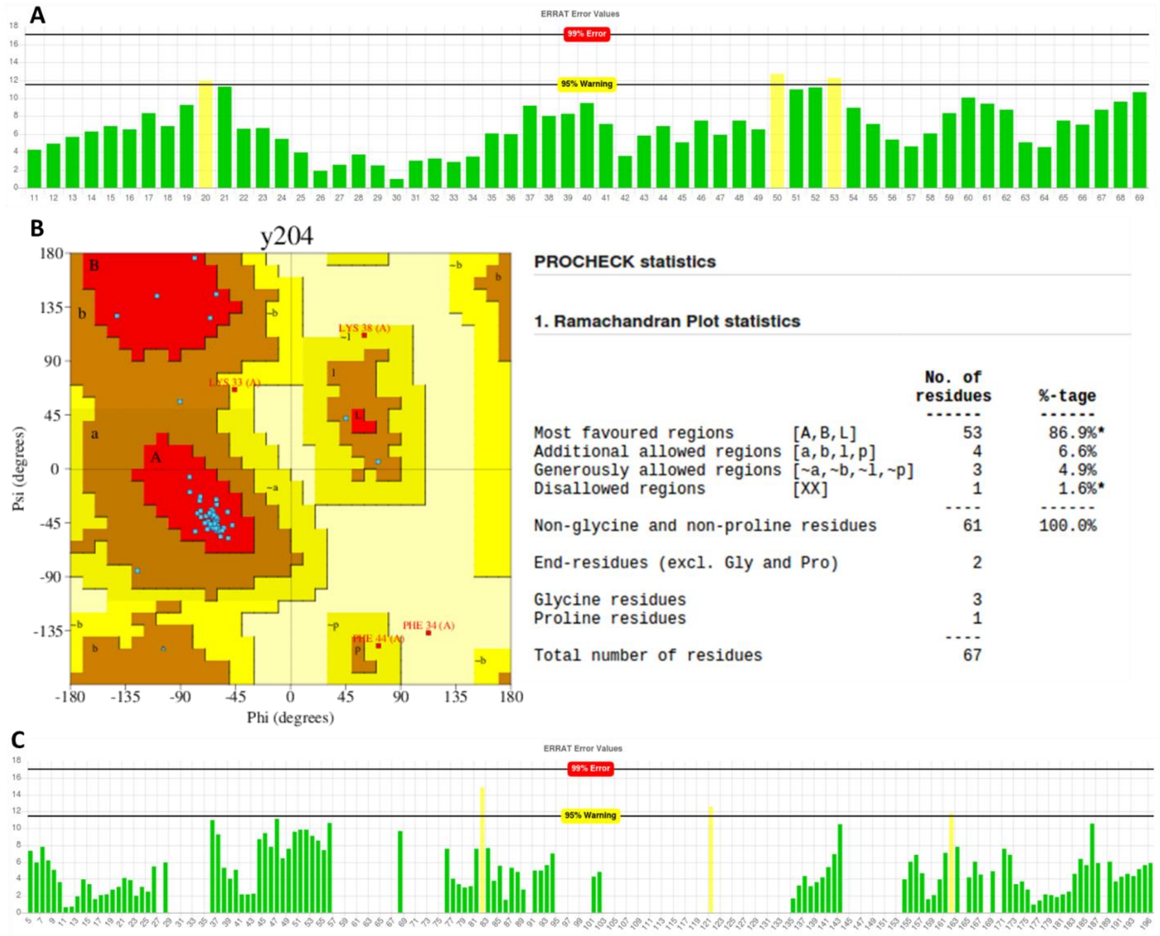


**Fig. S3.** Model quality assessment parameters for the models of residues 7-73 and 79-279; **(A)** ERRAT error plot for residues 7-73 (67 residues). % Error for each residue is plotted. Residues plotted in red colour indicate erroneous non-bonded interaction profile, hence unreliable region of the protein. **(B)** Ramachandran plot analysis of the model for residues 7-73. **(C)** ERRAT error plot for residues 80-276 (200 residues).

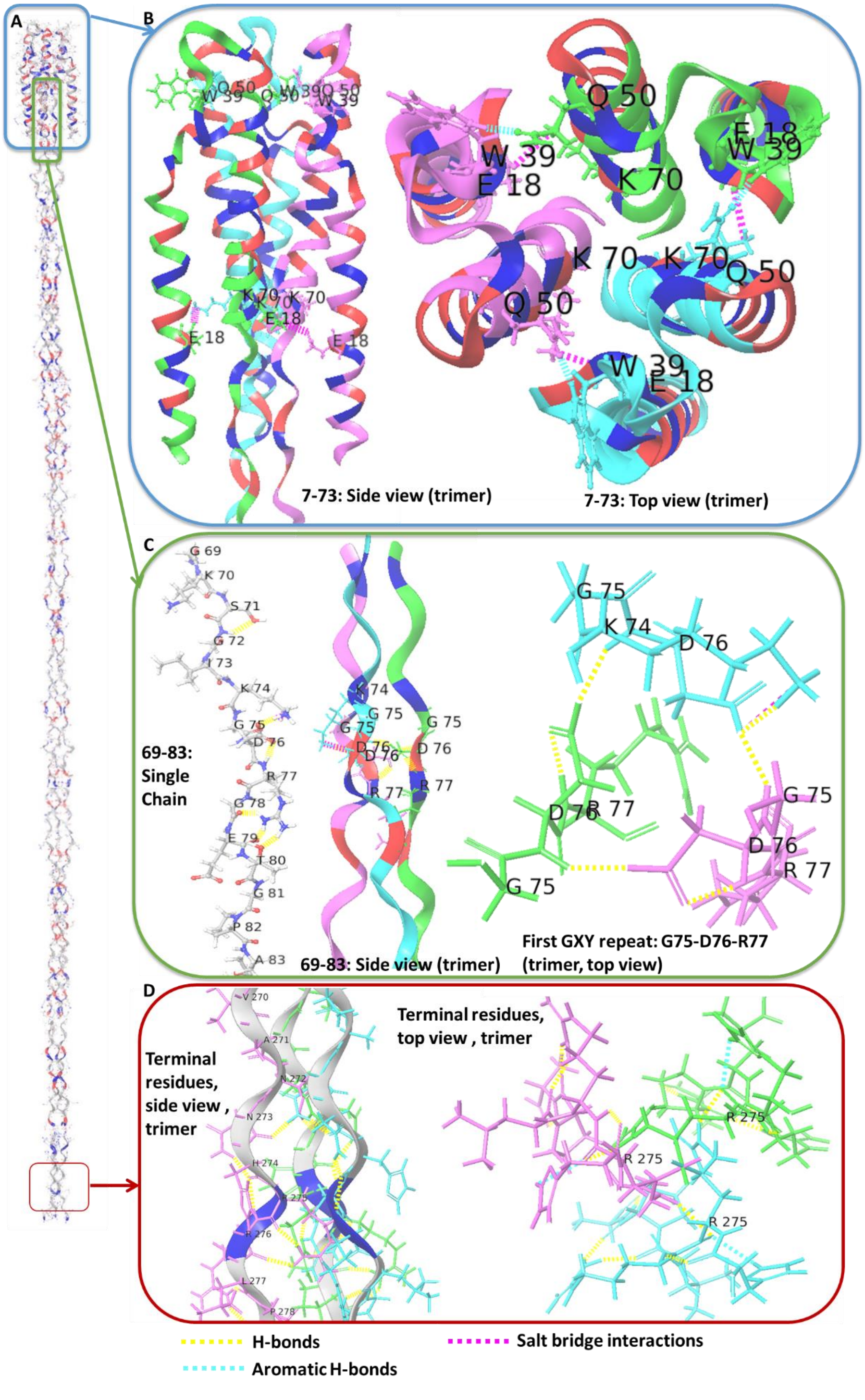


**Fig. S4. (A)** Scl1 trimeric model **(B)** N-terminal globular region, with inter chain interactions, **(C)** Inter chain and intra-chain non covalent interactions in the GXY regions (G75-R77), **(D)** Inter chain and intra-chain non covalent interactions in the C-terminal region (H274-R276).

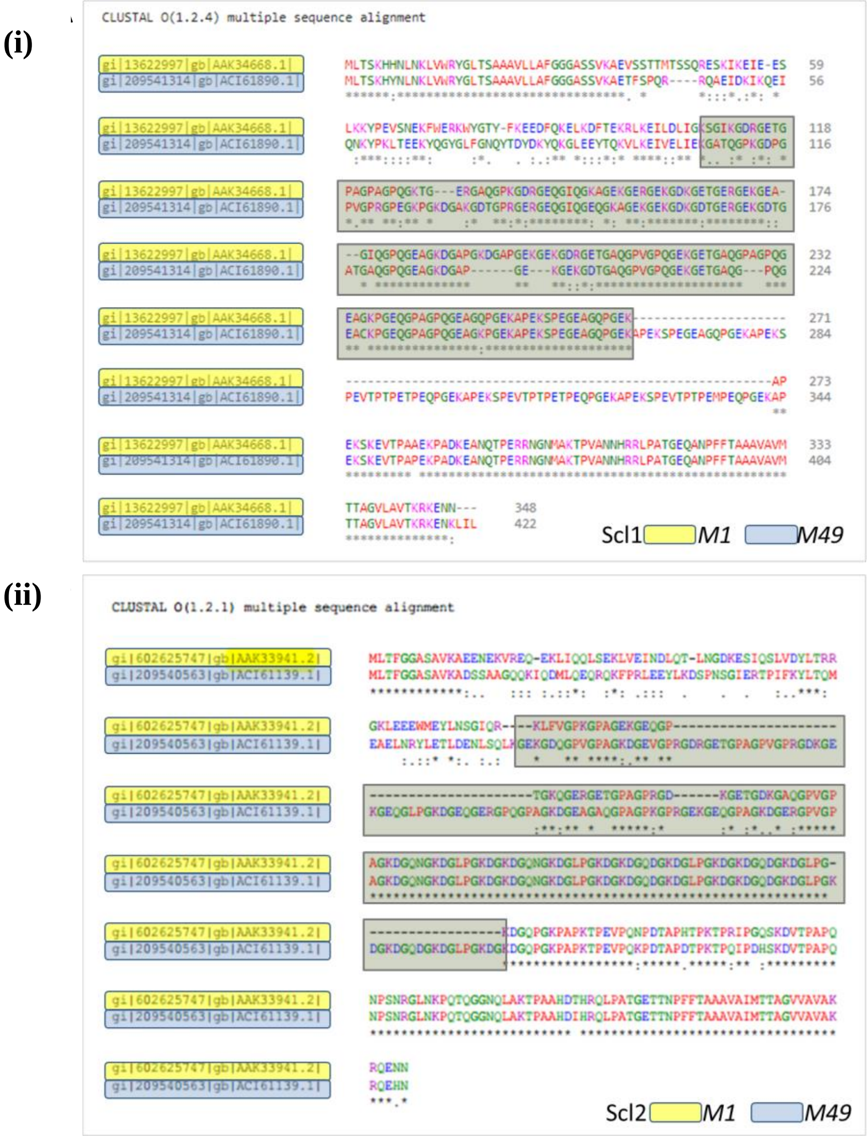


**Fig. S5A.** Similarity between major annotated sequences of GAS M1; M49, **(i) Scl1 proteins**, **(ii) Scl2 proteins.** The only difference is evidently seen in the CL-Domain lengths depicted green shaded box covering the CL-domain in each protein sequences.

(i)

(ii)

(iii)

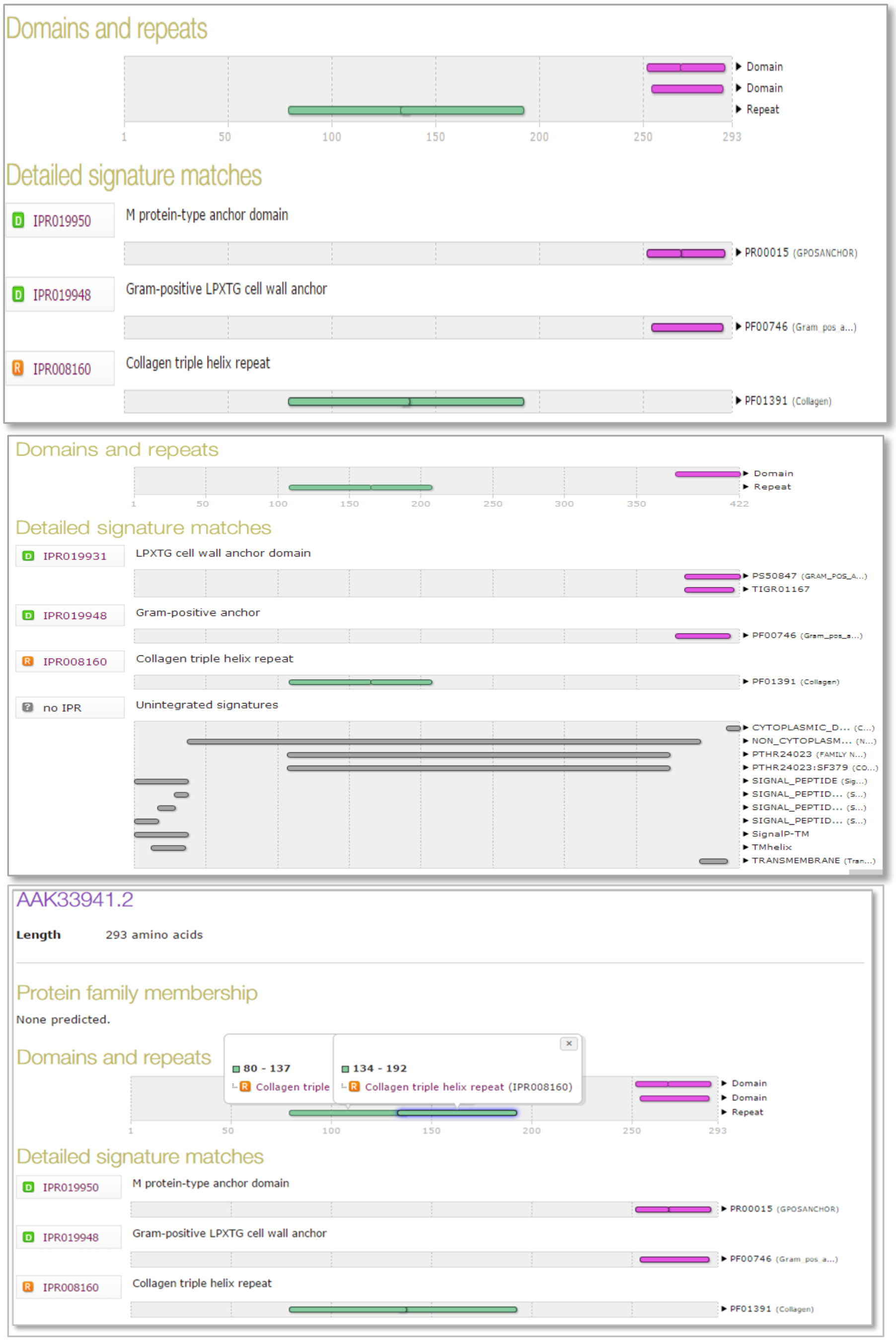


**Fig. S5B.** GAS Scl1 Protein-Domains Analysis, depicting the conserved protein-families from the database. **(i) GAS M1 Scl1. (ii) GAS M49 Scl1. (iii)** Collagen-like protein domain retrieved based on Protein family-database.

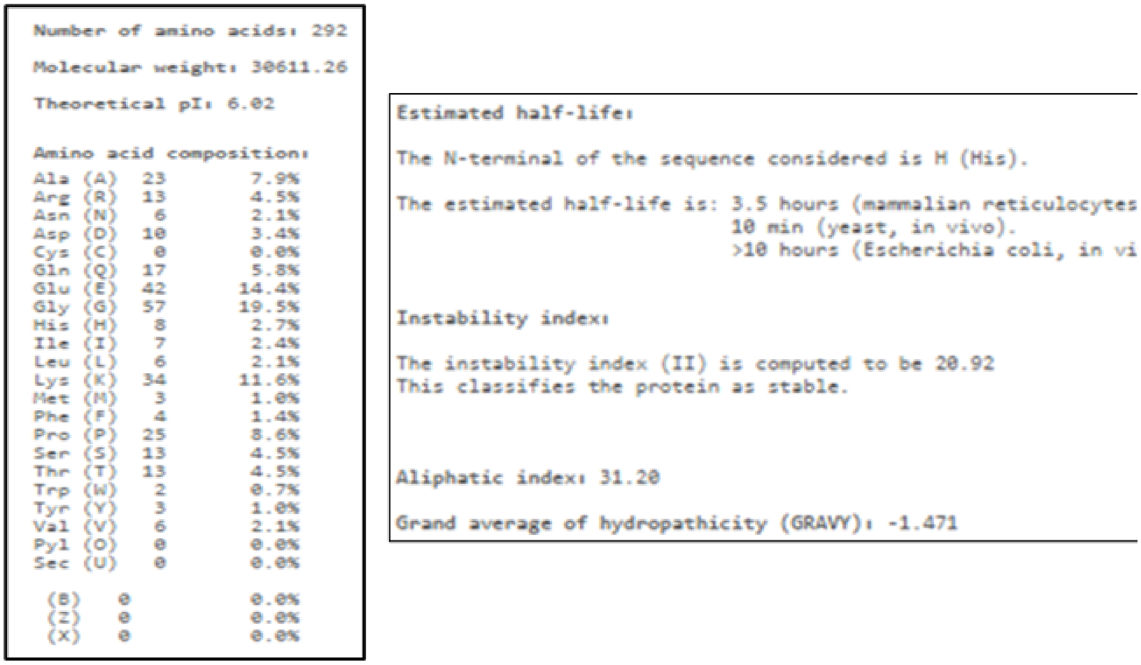

```
Number of amino acids: 292

Molecular weight: 30611.26

Theoretical pI: 6.02

Amino acid composition:
Ala (A)  23     7.9%
Arg (R)  13     4.5%
Asn (N)   6     2.1%
Asp (D)  10     3.4%
Cys (C)   0     0.0%
Gln (Q)  17     5.8%
Glu (E)  42    14.4%
Gly (G)  57    19.5%
His (H)   8     2.7%
Ile (I)   7     2.4%
Leu (L)   6     2.1%
Lys (K)  34    11.6%
Met (M)   3     1.0%
Phe (F)   4     1.4%
Pro (P)  25     8.6%
Ser (S)  13     4.5%
Thr (T)  13     4.5%
Trp (W)   2     0.7%
Tyr (Y)   3     1.0%
Val (V)   6     2.1%
Pyl (O)   0     0.0%
Sec (U)   0     0.0%

 (B)   0        0.0%
 (Z)   0        0.0%
 (X)   0        0.0%
```

```
Estimated half-life:

The N-terminal of the sequence considered is H (His).

The estimated half-life is: 3.5 hours (mammalian reticulocytes
                            10 min (yeast, in vivo).
                            >10 hours (Escherichia coli, in vi


Instability index:

The instability index (II) is computed to be 20.92
This classifies the protein as stable.


Aliphatic index: 31.20

Grand average of hydropathicity (GRAVY): -1.471
```

**Fig. S6A.** Analysis through Potparam/Expasy tool showing the meagre proportions of the conjugate amino acid residues (https://web.expasy.org/cgi-bin/protparam/protparam) and amino acid residue-proportion in pRS-1 backbone that predicted pRS-1-construct stable, at different conditions *viz.*, pH and expressed-protein stability in *E. coli*.

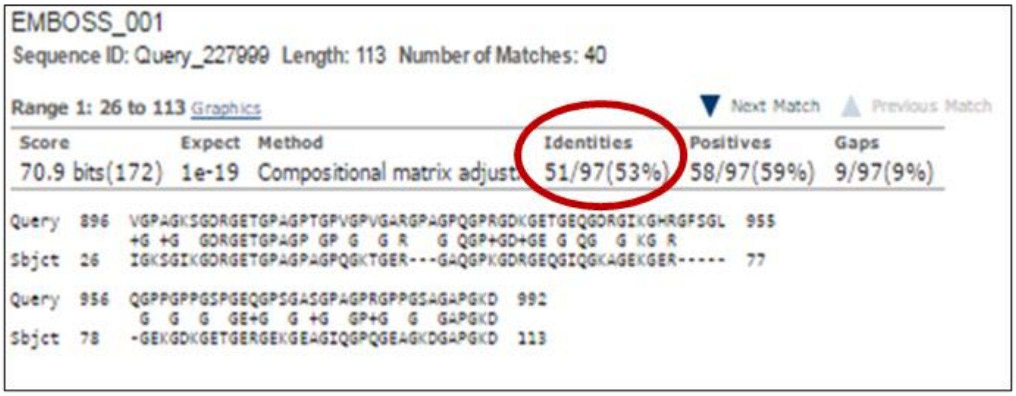

```
EMBOSS_001
Sequence ID: Query_227999  Length: 113  Number of Matches: 40

Range 1: 26 to 113 Graphics                         Next Match   Previous Match
Score            Expect  Method                          Identities   Positives    Gaps
70.9 bits(172)   1e-19   Compositional matrix adjust.    51/97(53%)   58/97(59%)   9/97(9%)

Query  896  VGPAGKSGDRGETGPAGPTGPVGPVGARGPAGPQGPRGDKGETGEQGDRGIKGHRGFSGL  955
            +G +G   GDRGETGPAGP GP G   G R   G QGP+GD+GE G QG   G KG R
Sbjct  26   IGKSGIKGDRGETGPAGPAGPQGKTGER---GAQGPKGDRGEQGIQGKAGEKGER-----  77

Query  956  QGPPGPPGSPGEQGPSGASGPAGPRGPPGSAGAPGKD  992
             G  G  G  GE+G  G +G  GP+G  G  GAPGKD
Sbjct  78   -GEKGDKGETGERGEKGEAGIQGPQGEAGKDGAPGKD  113
```

**Fig. S6B.** BLAST homology analysis for the rScl1-only collagen-like domain (CL) depicted 50% homology to human Collagen alpha-1(I) chain [position 162-1218 without the N- and C-terminal propeptide domains].

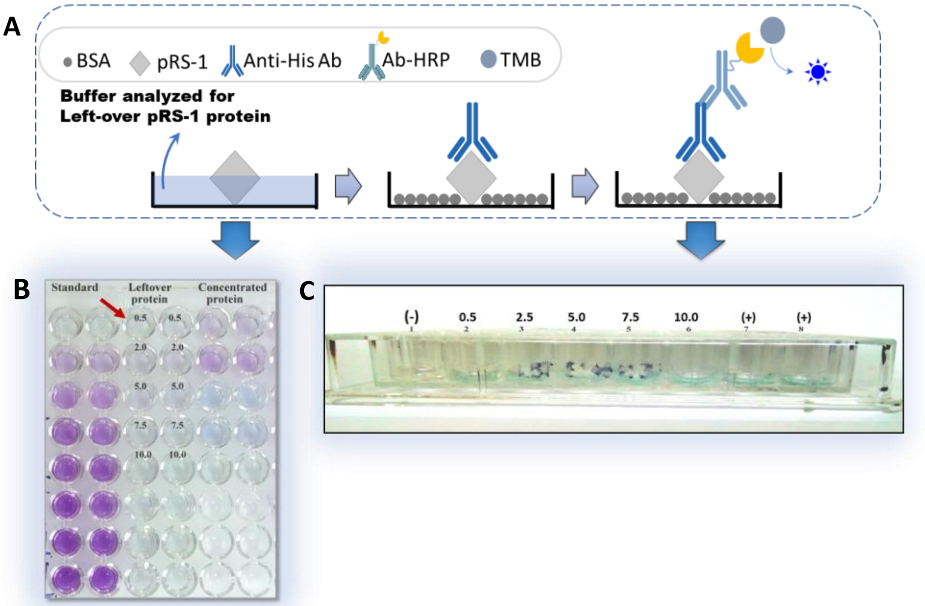


**Fig. S7. (A)** Schematic depicting pRS-1 matrix-like coating property in 96 well-plate, pRS-1 (Bicarbonate carbonate buffer) coated, blocked and detected by Anti-His primary antibody, **(B)** Plate assay showing estimation of leftover pRS-1 in coated-plates by BCA. First 2 lanes: BCA standard; middle 2 Lanes: Leftover buffer (Arrow in red); Last 2 lanes: pRS-1 protein, **(C)** pRS-1 colourimetric detection (Blue). Well: Control without pRS-1 (1); Coated pRS-1 in 0.3 M bicarbonate carbonate buffer (2-6), showing blue colourimetric reaction; Coated pRS-1 in 300 mM Imidazole Elution buffer (7, 8).

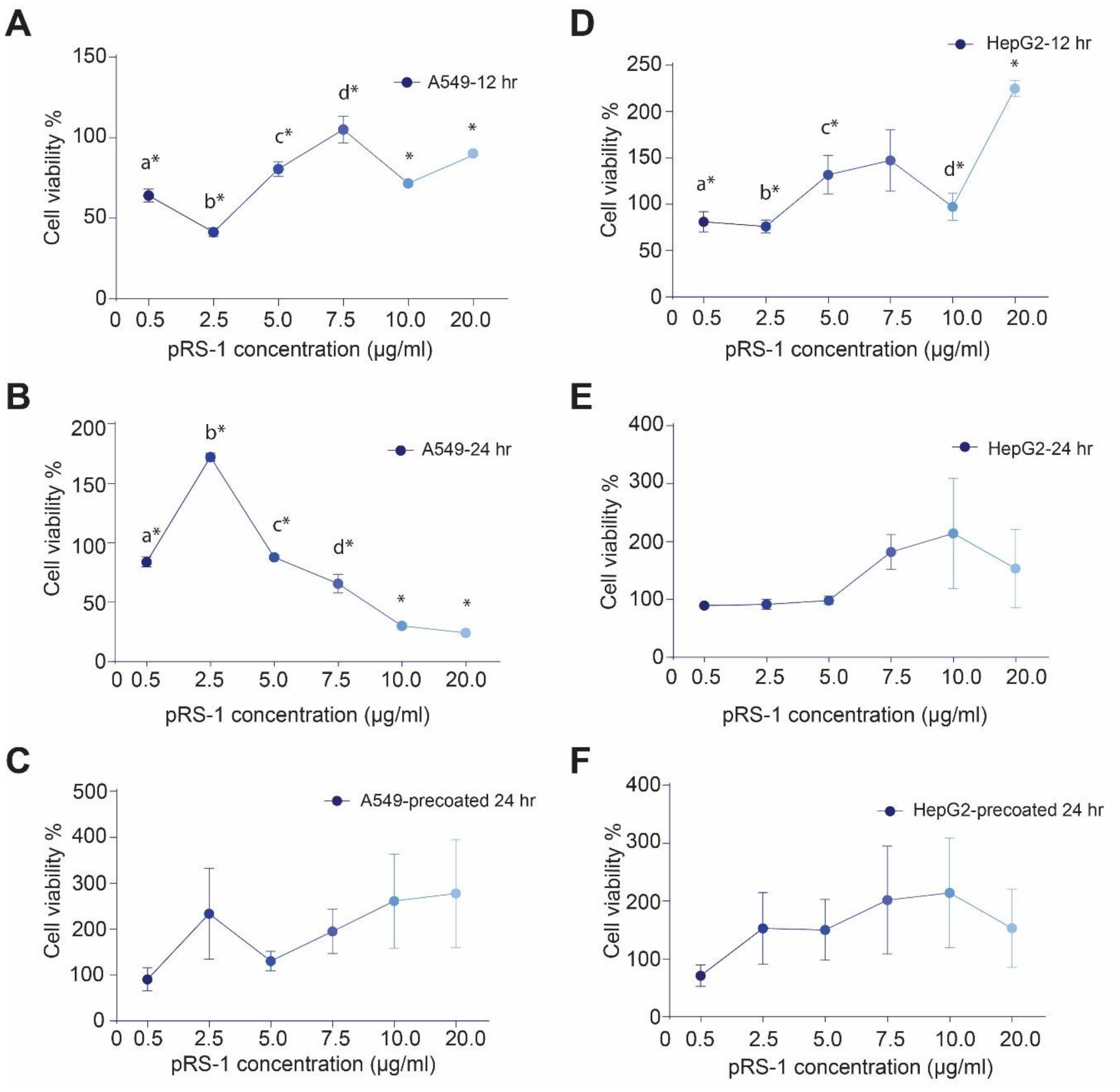


**Fig. S8.** Cell-Cytotoxicity and Cell-Viability analysis. A549 cells; **(A)** 12 hrs, **(B)** 24 hrs, **(C)** 24 hrs stored precoated plate study; HepG2 cells; **(D)** 12 hrs, **(E)** 24 hrs, **(F)** 24 hrs stored precoated plate study.

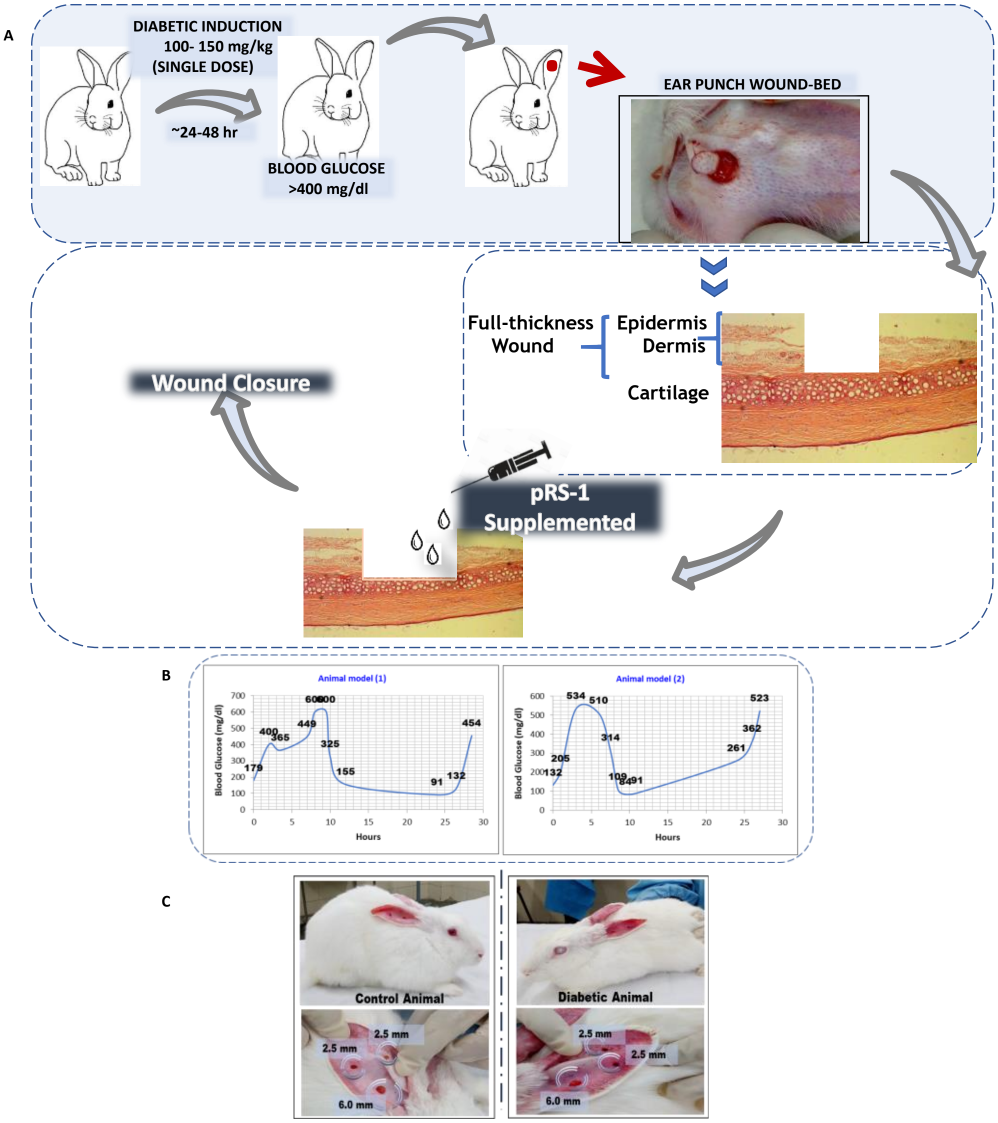


**Fig. S9. (A)** Schematic showing the study workflow showing experimental-diabetic model generation within 24 hrs of Alloxan administration in New Zealand White Rabbits and full-thickness cutaneous wound model. **(B)** *In vivo* diabetic-induction confirmation through Blood glucose response **(C)** Control and Diabetic rabbit depicted with 2.5 mm and 6.0 mm wound models.

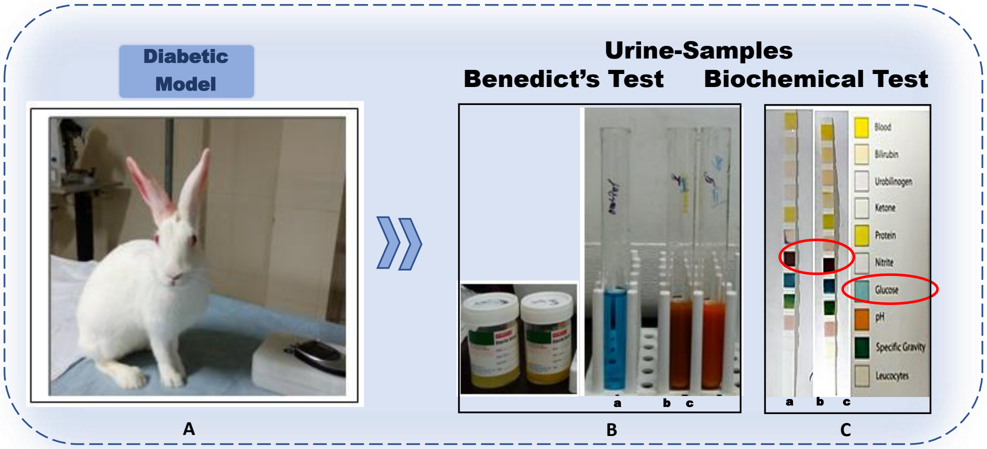


**Fig. S10.** Biochemical confirmation of the developed and sustained experimental-diabetic (Alloxan) model in New Zealand White rabbit after few months. **(A)** Diabetic model. Urine Samples from the few months diabetic rabbit models, **(B)** Benedict's quantitative test for the reducing sugars depicting >400mg/dl. **(C)** Biochemical Urine-strips **(a, b)** Urine-test strips of the diabetic-models, with deep-red (++++) colour-indicator for quantitative estimation of high-glucose in urine, **(c)** Control-strip with unchanged blue colour for glucose.

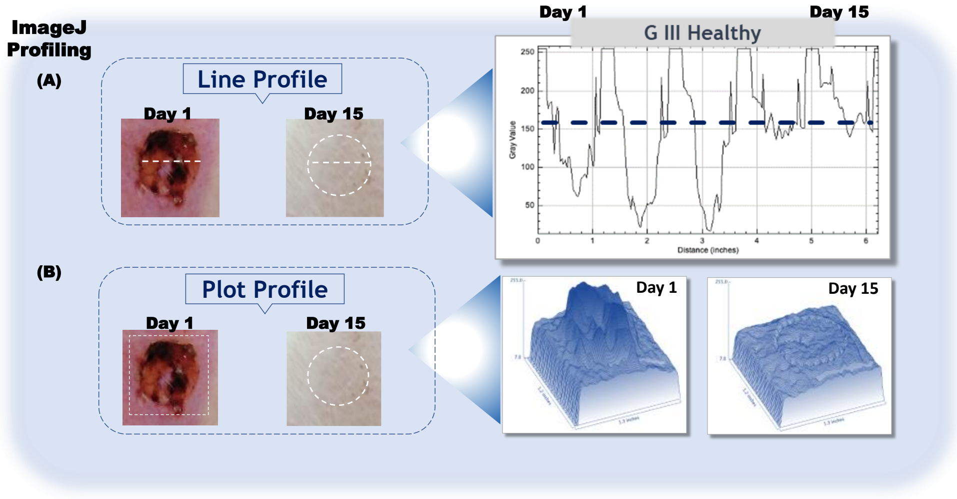


**Fig. S11.** ImageJ morphometry analysis using the Line & Plot profile tools. (**A)** Line profile with baseline (indicated in blue dotted line) selection made by including the surrounding normal-tissue in the wound-bed analysis through day 1 to day 15 showing the graphical output as the representative profiles for an entire group (Group-wise analysis). **(B)** Plot profile shows contoured-texture represented to show morphometric-estimation of depth and intensity of wound-bed comparative to the baseline surrounding tissue (indicated by the white dotted-square in the day 1 wound-bed) and graphical profile clearly defines comparative depth of initial wounds (day 1) and a complete closure at day 15. This method was devised to study variation in intensities and record quantitative morphometry of wound-beds.

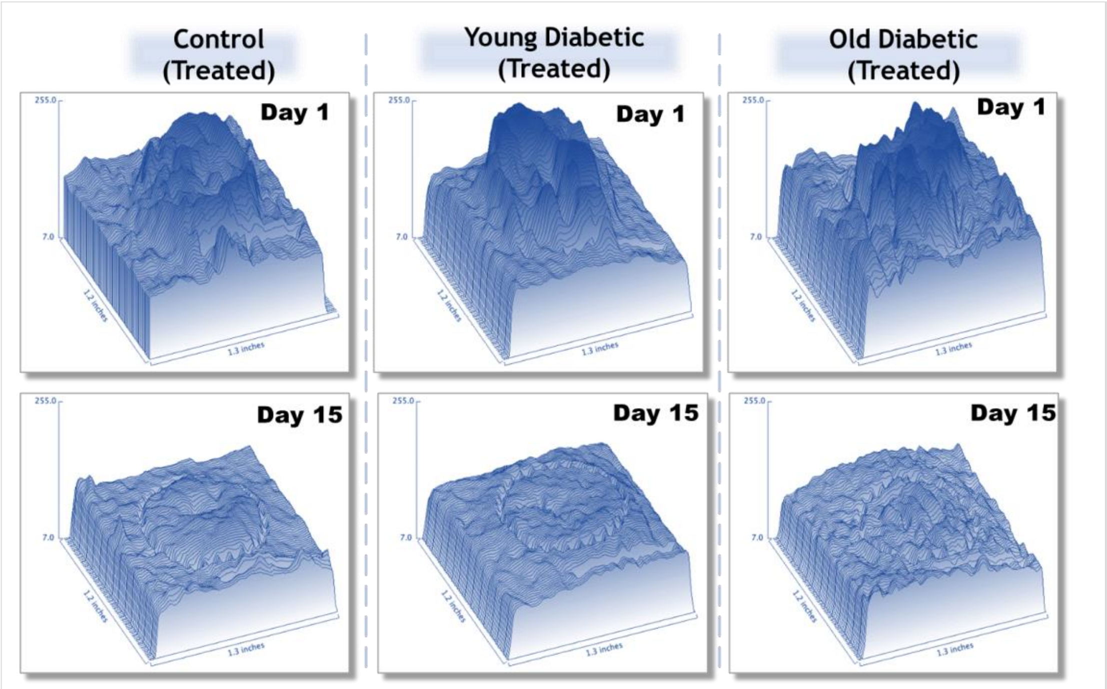


**Fig. S12.** Intensity Plot profiles for the Old-diabetic group. ImageJ software analysis tools were employed for morphometric-estimation of pRS-1 treated wound-beds in Control; Young-diabetic & Old-diabetic models after 15 days of wound-induction. Upper panels show the baseline normal-tissue along with the raised contours indicating Day 1 wound-area plotted in sync to their intensity of depth and texture thus, measuring their patterns morphometrically and the same can be evidently seen in lower panel showing Day 15, in all the pRS-1 treated groups, where circular wound-bed edges are indicative of the baseline (drawn in the figures to mark the healed region) showing the intensity of the healed wound-bed levelled to that of the adjacent normal-tissue.

A

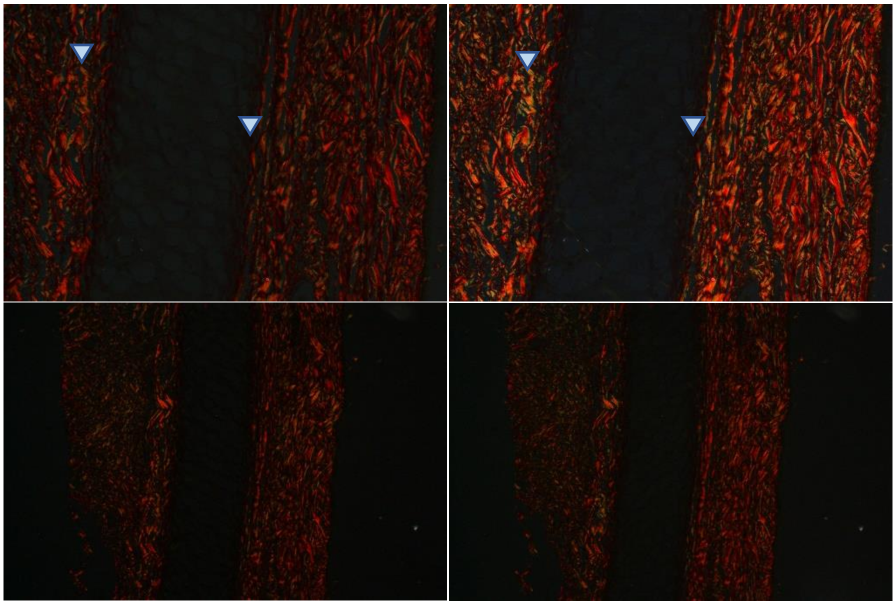

B

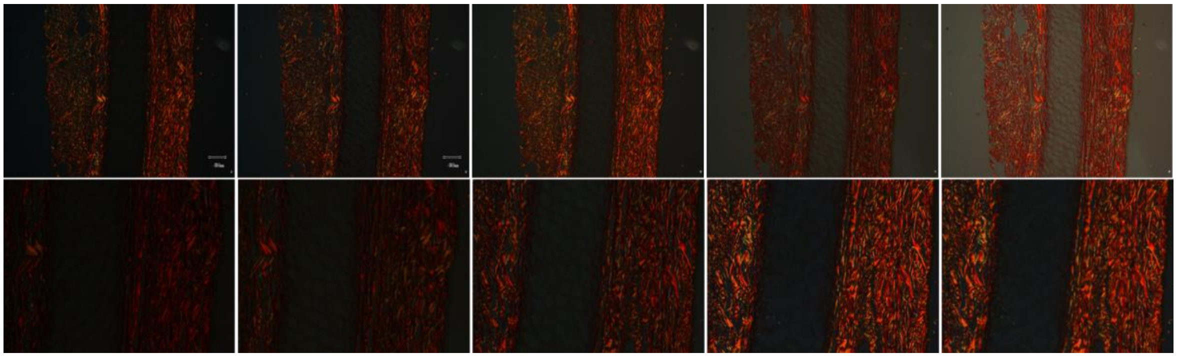

C

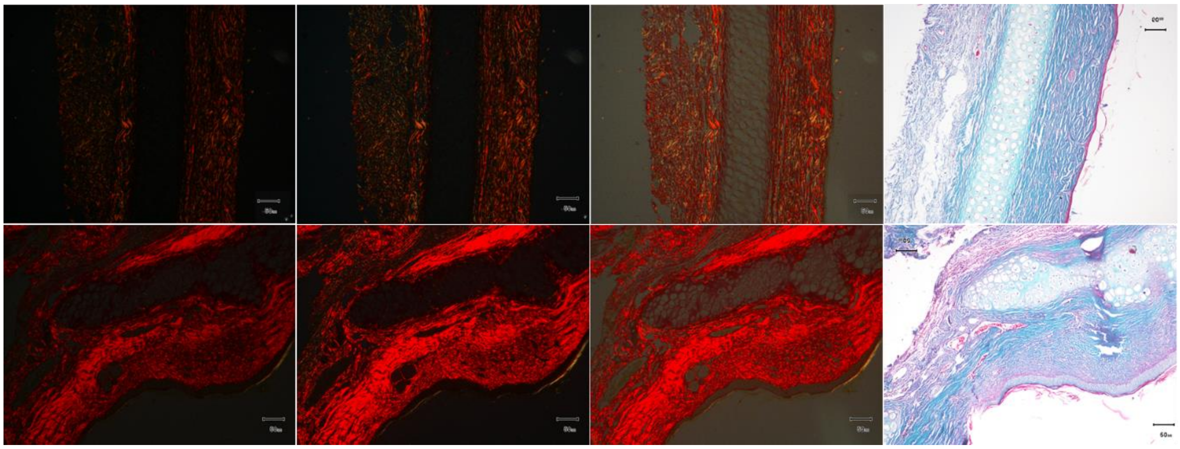

**Fig. S13.** Picrosirius red stained cutaneous full-thickness, rabbit-ear sections depicting the birefringence. (**A)** Plane polarized images in Healthy-cutaneous tissue showing the green-yellow birefringent collagen type III fibers denoted in Blue arrow-heads (Original magnification 200x upper panel, 100x lower panel), (**B)** Detailed panel showing the plane polarized images of the same section as shown in (**A)**, **(C)** Picro Sirius Red stained sections (Birefringence): Quantitation of collagen-content in the Control-healthy and >6 months Diabetic-model cutaneous-tissues; distinct green-yellow, thin, collagen type III fibers in Control-sections as against the red, coarsely stained, thick, collagen type I bundles in Old-Diabetic models. Upper panel-Control-sections; Lower panel- >6 months Diabetic-sections; last column in both panels is Gomori trichome sections; scale bar is 50 µm.

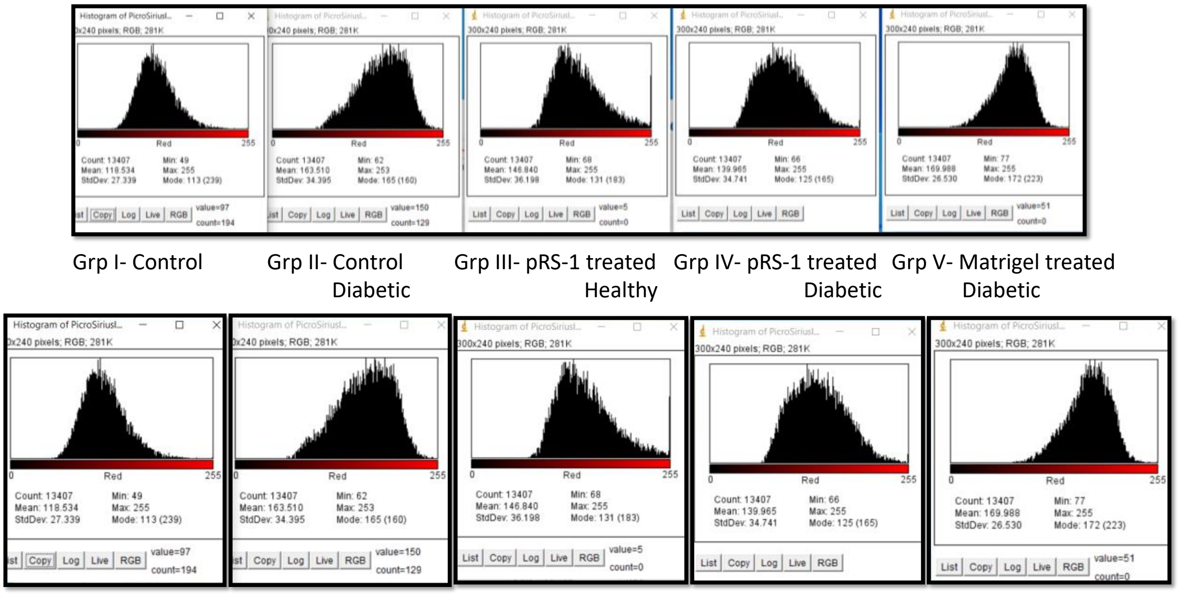


**Fig. S14.** ImageJ analysis of the Picro Sirius red stained sections in Figure 7A showing the red intensity histograms from each section of age-matched rabbit models (upper panel) and zoomed image (lower panel); from left, **Group I (118)-** Untreated Control Healthy; **Group II (163)-** Untreated Control Diabetic; **Group III (146)-** pRS-1 Treated Healthy; **Group IV (139)-** pRS-1 Treated Diabetic; **Group V (169)-** Matrigel treated Diabetic. The histograms are shown in Figure 7B in a vertical alignment to compare the red intensity which is restored in pRS-1 treated Diabetic- Group IV and is lesser than both Control Diabetic-Group II and Matrigel treated Diabetic- Group V indicating the healthy collagen-proportion as estimated through Picro Sirius quantitation.

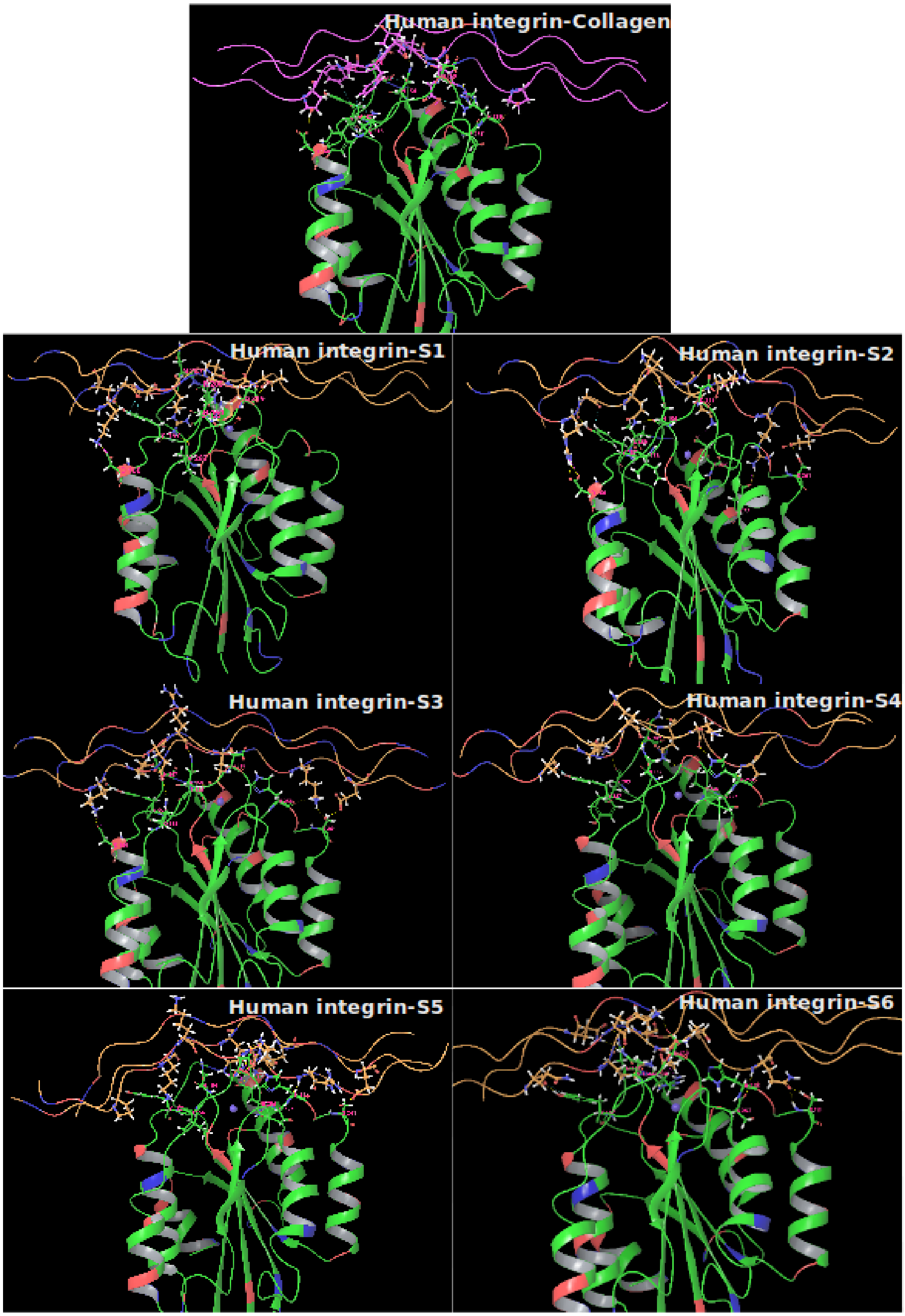

Human integrin-Collagen
Human integrin-S1
Human integrin-S2
Human integrin-S3
Human integrin-S4
Human integrin-S5
Human integrin-S6

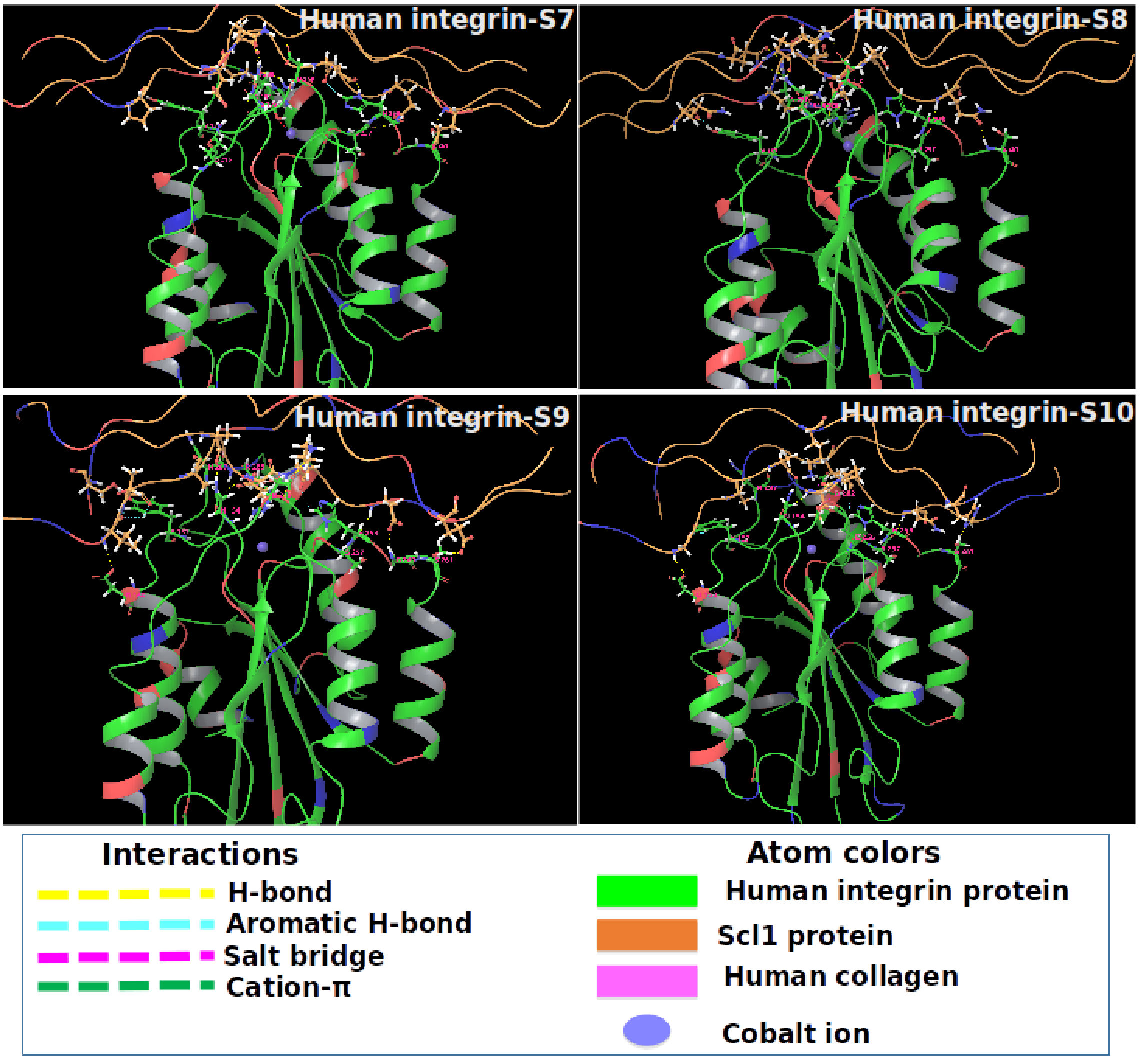


**Fig. S15.** Complexes of human integrin with Scl1 peptides and collagen

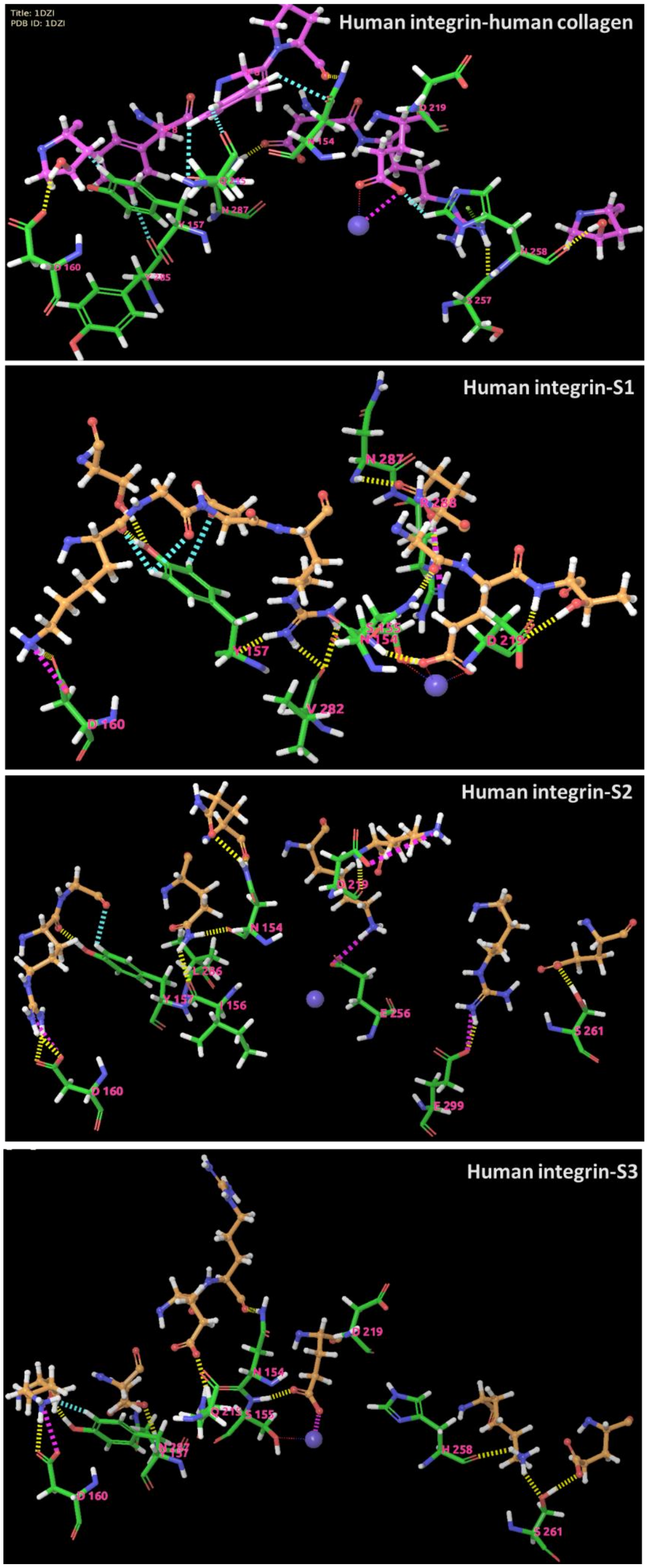
Title: 1DZI
PDB ID: 1DZI
Human integrin-human collagen
D 219
N 154
N 287
N 157
D 160
H 258
S 257
Human integrin-S1
N 287
N 157
N 154
D 219
V 282
D 160
Human integrin-S2
N 154
N 157
I 156
E 256
S 261
D 160
Human integrin-S3
D 219
N 154
S 155
H 258
D 160
S 261

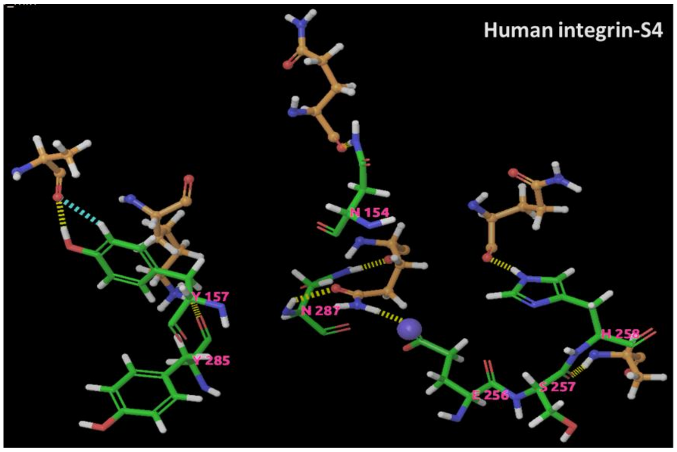
Human integrin-S4
N 154
N 287

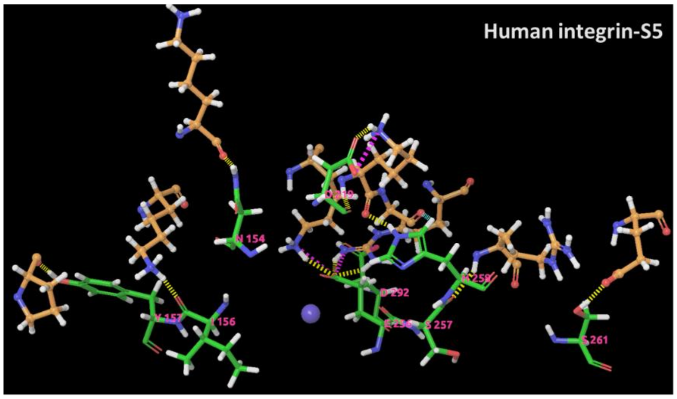
Human integrin-S5

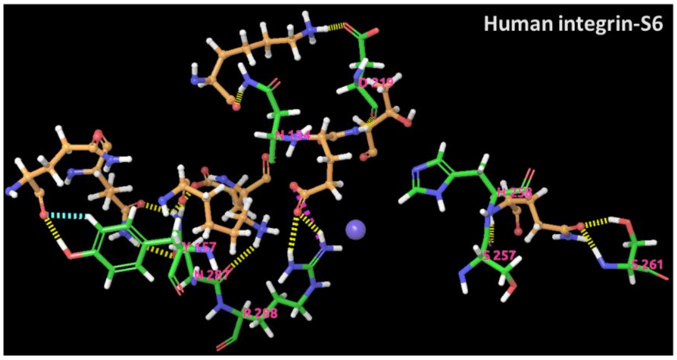
Human integrin-S6

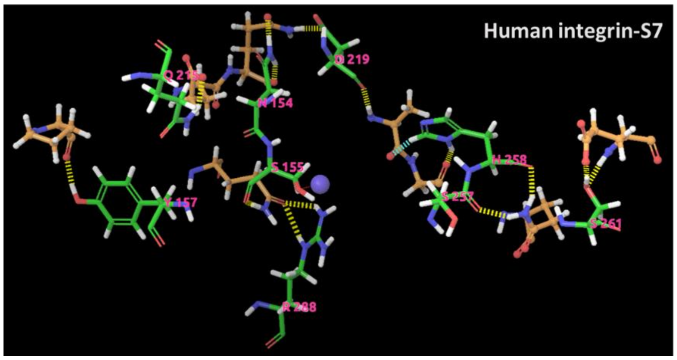
Human integrin-S7
N 154

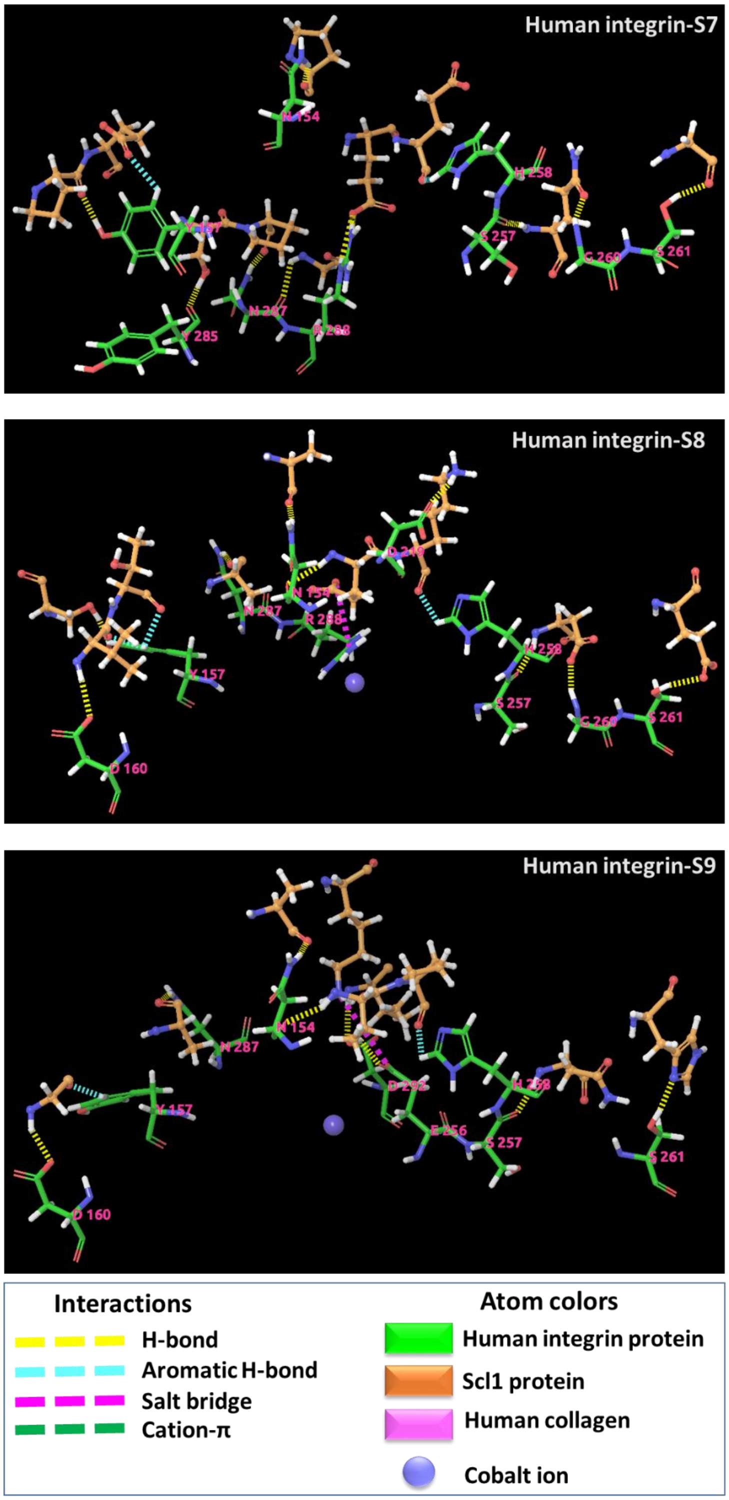


**Fig. S16.** Non covalent interactions between human integrin and Scl1 trimeric peptides/human collagen.
